# Gq-mediated calcium dynamics and membrane tension modulate neurite plasticity

**DOI:** 10.1101/661975

**Authors:** Katherine M. Pearce, Miriam Bell, Will H. Linthicum, Qi Wen, Jagan Srinivasan, Padmini Rangamani, Suzanne Scarlata

**Author notes:** both these authors contributed equally to this study (Katherine M. Pearce, Miriam Bell).

## Abstract

The formation and disruption of synaptic connections during development is a fundamental step in neural circuit formation. Subneuronal structures such as neurites are known to be sensitive to the level of spontaneous neuronal activity but the specifics of how neurotransmitter-induced calcium activity regulates neurite homeostasis are not yet fully understood. In response to stimulation by neurotransmitters such as acetylcholine, calcium responses in cells are mediated the Gαq/phospholipase Cβ (PLCβ)/ phosphatidylinositol 4,5 bisphosphate (PI(4, 5)P_2_) signaling pathway. Here, we show that prolonged Gαq stimulation results in the retraction of neurites in PC12 cells and rupture of neuronal synapses by modulating membrane tension. To understand the underlying cause, we dissected the behavior of individual components of the Gαq/PLCβ/PI(4, 5)P_2_ pathway during retraction, and correlated these to the retraction of the membrane and cytoskeletal elements impacted by calcium signaling. We developed a mathematical model that combines biochemical signaling with membrane tension and cytoskeletal mechanics, to show how signaling events are coupled to retraction velocity, membrane tension and actin dynamics. The coupling between calcium and neurite retraction is shown to be operative in the *C. elegans* nervous system. This study uncovers a novel mechanochemical connection between the Gαq/PLCβ/(PI(4, 5)P_2_ pathway that couples calcium responses with neural plasticity.

## INTRODUCTION

Throughout the course of an organism’s life, different neuronal connections break and reform to generate new electrical patterns that allow for optimal function during development and through adulthood. This plasticity of neuronal connections allows the rewiring of circuitry necessary for memory and learning (e.g.(1) (2) (3)). Understanding the factors that permit appropriate and efficient rewiring is essential for understanding both developmental and neurodegenerative diseases.

Calcium is a key mediator of neuronal functions such as axonal growth, neurite protrusion, and spinogenesis (4, 5). Previous studies have shown that spontaneous activity in neurons can results in a frequency dependent rate of axon elongation such that the rate of axon elongation is inversely proportional to the frequency of calcium transients (6, 7). The role of calcium in neurite growth and protrusions associated with development has been studied in different contexts (e.g (8)). This outgrowth process is stimulated by neurotrophic factors that carefully regulate the spatiotemporal aspects of intracellular calcium. Neurite growth can also be triggered by stress, pharmacological agents, or starvation (9, 10). These factors may occur during disease states and contribute to inappropriate neurite growth, which can result in an increased number of smaller and non-productive neurites, as seen in autism (10).

Neurodegenerative diseases are associated with the disruption of normal calcium signaling. Specifically, diseases such as autism and amyotrophic lateral sclerosis (ALS) have been linked to over-excitation of neurons or excitotoxicity caused by abnormal calcium homeostasis (11). However, the link between calcium homeostasis and the mechanical processes underlying neurite retraction remain poorly understood. Understanding the link between dysfunction of cellular calcium, especially prolonged signaling, and neuronal function is critical to understand the mechanisms that underlie diseases typified by inappropriate neuronal excitation. While previous research has focused on the short-term effects of elevated cellular calcium after stimulation and its effects on neurite growth or on spontaneous spiking activity (6, 7, 12), little is known about the effect of extended elevated calcium on these same neurites over time, i.e. cases that mimic an extended overstimulated state. In this study, we investigate the effects of prolonged stimulation of Gαq/PLCβ/PIP_2_/calcium signaling pathway by the neurotransmitter acetylcholine, in a model neuronal cell line, PC12, and in the neuronal network in a small organism, *Caenorhabditis elegans*.

Calcium signals in neurites can be generated by several mechanisms. In synapses, postsynaptic cells receive an influx of calcium directly through the opening of transmembrane channels in response to neurotransmitter release (13). Neurotransmitters, such as acetylcholine, activate G protein coupled receptors (GPCRs) on the plasma membrane to generate the release of calcium from intracellular stores (9). Furthermore, neurotropic factors activate receptor tyrosine kinases to increase intracellular calcium through another phospholipase C family, PLCγ. Downstream from these events are the opening of calcium-activated calcium channels on the endoplasmic reticulum (ER) membrane. Many of these channels are thought to respond to the increase in cytosolic calcium and to changes to the physical properties of the plasma membrane. These increases in cellular calcium regulate specific transcription factors and post-transcriptional processes that lead to appropriate downstream responses (14).

In this study, we show that neurites will completely retract to the soma by extended stimulation of the GPCR/Gαq/PLCβ pathway. This pathway mediates signals from many hormones and neurotransmitters such as acetylcholine, dopamine, histamine, and melatonin (15). Signaling begins with ligand binding to its specific G protein coupled receptor to activate the Gαq family of G proteins by exchange of GTP for GDP. The GTP-bound Gαq subunits then actives phospholipase Cβ (PLCβ). PLCβ catalyzes the hydrolysis of phosphatidylinositol 4,5-biphosphate (PIP2) to inositol 1,4,5-triphosphate (IP3) and diacylglycerol (DAG). IP_3_ binds to receptors on the endoplasmic reticulum allowing the release of calcium from intracellular stores into the cytoplasm. The elevated calcium can then change the activity of a variety of intracellular proteins to generate a specific cell response.

Incorporated in this pathway are a number of positive and negative feedback loops. The initial calcium release generated by PLCβ in turn stimulates the highly active PLCδ that synergizes the calcium response (16). Increased calcium also opens calcium-induced calcium channels, further strengthening calcium responses (17). Negative feedback comes from the internalization of ligand-bound receptors into endosomes over a few minutes that typically return unbound receptors to the plasma membrane over a period of 20-30 min (18). Additionally, Gαq has a GTPase activity that returns it to the basal state and this activity is stimulated by the GTPase activating (GAP) activity of PLCβ (19), along with RGS proteins that quickly turn-off the signal (20). It is notable that these feedback loops are highly sensitive to the concentration of the pathway components, the concentrations of competing species as well as the physical state of the plasma membrane which drives aggregation of the ligand-bound receptor and internalization, the accessibility of (PI(4, 5)P_2_ substrates to PLCβ and PLCδ, and the opening of calcium channels.

Components of the Gαq/PLCβ/PI(4,5)P_2_ pathway are also involved in the regulation of membrane tension and cytoskeletal remodeling. The coupling between membrane tension and cytoskeletal adhesion through (PI(4, 5)P_2_ was established nearly two decades ago by Sheetz and coworkers (21, 22). In a series of elegant experiments, they showed that PIP_2_ regulates cytoskeleton-plasma membrane adhesion (22) and that membrane tension plays a critical role in actin remodeling, membrane trafficking, and cell motility (23–25). Subsequently, researchers have continued to establish and identify increasing roles played by the plasma membrane tension and cortical tension in different cellular reorganization processes (26) (27, 28). Specifically, in neurons, dynamics of growth cone formation, neurite protrusion, and axonal contractility have been shown to be mechanochemically coupled processes (29, 30).

Despite these advances in neuronal biophysics, our understanding of synaptic rupture and neurite retraction remains incomplete especially in terms of the mechanisms that underlie overexcitatory responses. In this study, we set out to investigate the effects of prolonged (i.e. several minutes) agonist stimulation of the Gαq signaling pathway in a model neuronal cell line (PC12). We find that prolonged stimulation results in tension-driven neurite retraction unlike the behavior seen in shorter exposures to neurotransmitters. To understand the factors that underlie the observed neurite retraction, we followed the individual components during the process and developed a mathematical model to predict the effects of calcium stimulation by acetylcholine (ACh) and its extended response on retraction through mobilization of the pathways that impact neurite retraction.

Our predictive models showing that membrane tension and actin reorganization are coupled to calcium dynamics through Gαq/PLCβ/(PI(4, 5)P_2_ were verified in cultured PC12 cells as well as in the neuronal network of the nematode *Caenorhabditis elegans.* The 302 neurons that (31)comprise the nervous system of *C. elegans* have been well characterized and these organisms have been used as models to understand neurite formation and retraction (31). Because of their optical clarity, *C. elegans* allows us to monitor the effects of acetylcholine stimulation on synapses in real time by microscopy. We find that the *C. elegans* neural architecture exhibits the same retraction behavior when exposed to Gαq agonists showing rupture along the spine in the nerve ring, suggesting that the coupling between membrane tension and calcium dynamics occur on the organism level. Taken together, our studies connect signaling processes with mechanical effects that allow us to predict the signaling conditions that shift from outgrowth and maintenance to retraction.

## 2. RESULTS

### Prolonged exposure to carbachol causes neurite retraction

Cultured PC12 cells differentiate to a neuronal phenotype upon treatment with nerve growth factor (NGF) which initiates activation through TrkA receptors (32). This treatment results in growth of neurites from the cell body that extend to roughly 3 times the length of the body over a 36-hr period. These neurites can then connect with neurites from other cells resulting in long tubular structures (33).

Addition of a Gαq agonist to the cells activates PLCβ to catalyze the hydrolysis of (PI(4, 5)P_2_. This hydrolysis releases Ins(1,4,5)P_3_ into the cytosol that then binds to Ins(1,4,5)P_3_ receptors on the ER to release calcium from intracellular stores. In addition to eliciting calcium signals, (PI(4, 5)P_2_ hydrolysis can exert mechanical effects on cells by altering membrane tension and actin-membrane adhesion (21) (22, 27). We sought to understand the interactions between the Gαq/PLCβ pathway and mechanical features of membrane-actin interaction (Fig. lA, B).

We found that when we add a Gαq agonist, such as carbachol or bradykinin, to PC12 cells, the well-formed neurites become thinner, their connections rupture (compare Fig lC to Fig. lE, S1) and they retract toward the soma after a period of 5-10 minutes (Fig. lE, F). After retraction, the excess membrane appears as blebs around the sides of the cell (Fig. lG). This behavior was seen in every cell viewed in over 100 experiments, but was not seen when a PLCβ inhibitor was added, or a Gαi agonist, such as isoproterenol was used. The initiation and rate of retraction depended on the particular treatment and condition of the experiment, as described below. The extent of retraction in a specific time period was robust to the length and thickness of the neurite.

**Figure 1:**
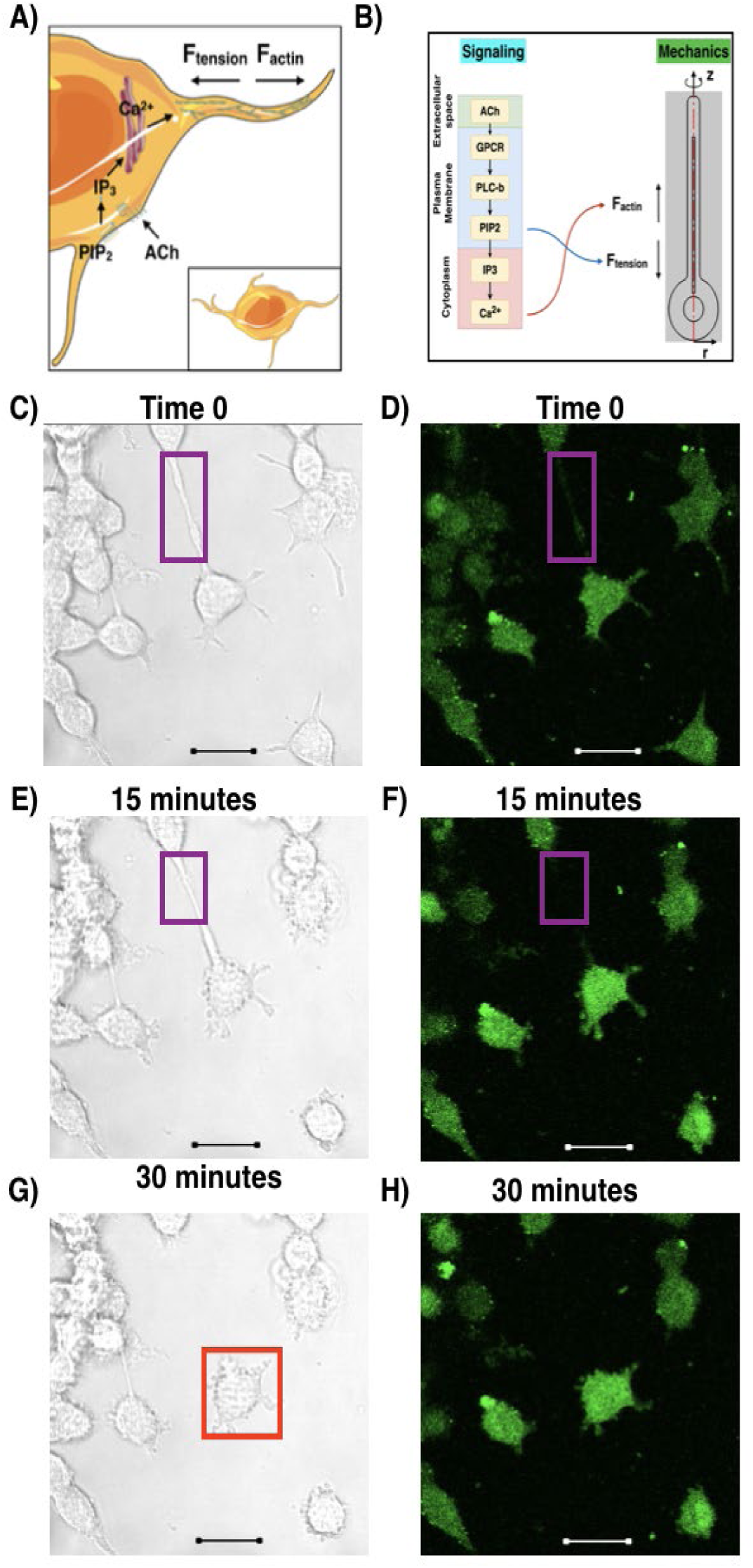
Neurite retraction in PC12 cells is induced by calcium stimulation. **A)** *Cartoon of the mechanochemical events underlying neurite retraction in a PC12 cell.* The figure depicts the interconnection between calcium signaling and force regulation due to membrane cortical tension and actin remodeling. **B)** *Schematic showing the* c*oupling between signaling and mechanical changes within the PC12 cell*. We identify membrane cortical tension and actin dynamics as key players controlling neurite retraction rates, and link calcium dynamics to actin reorganization and force generation, and PI(4,5)P_2_ hydrolysis to membrane tension change and tension force generation. Using this coupled model, we are able to reproduce the neurite retraction behavior observed experimentally. **C-D)** *Sample images of differentiated PC12 cells before stimulation*. Confocal phase contrast (C) and fluorescence images of cells loaded with a fluorescent calcium sensor, Calcium Green, are shown (D). (E-G) *Stimulation of cells with carbachol results in neurite thinning, retraction and synaptic rupture*. 15 minutes after the addition of carbachol, we observe membrane retraction (e.g. purple box), and thinning of the neurite (E-F). 30 minutes after carbachol addition, we see a complete retraction of the neurites into the soma and membrane blebbing at the retraction sties (e.g. red box) (G-H). In all images, the scale bar is 20µm. Identical behavior was seen in 20 cells.

### Neurite retraction is coupled with Gαq/PLCβ/calcium pathway

We followed the activation of the Gαq/PLCβ/calcium signaling pathway upon stimulation in single PC12 cells using the fluorescent calcium indicator, Calcium Green, to determine whether neurite retraction is concurrent with activation. Our approach was to follow some of the molecular constituents of the Gαq pathway and determine the temporal correlation between activation of the individual signal components and neurite retraction. The retraction velocities of a generic Gαq-coupled receptor (i.e. the bradykinin receptor type 2 or B2R) were measured by transfecting cells with a fluorescent-tagged construct and stimulating the cells with agonist (i.e. bradykinin). We note that B2R is not endogenous to PC12 cells, thus allowing us to compare Gαq-associated retraction with those that result from stimulation of endogenous muscarinic receptors. We find identical retraction behavior for neurite retraction after stimulating the B2R transfected cells with bradykinin as those that result from carbachol stimulation (Fig. 2). These results support a connection between neurite retraction and Gαq/PLCβ activation.

**Figure 2.**
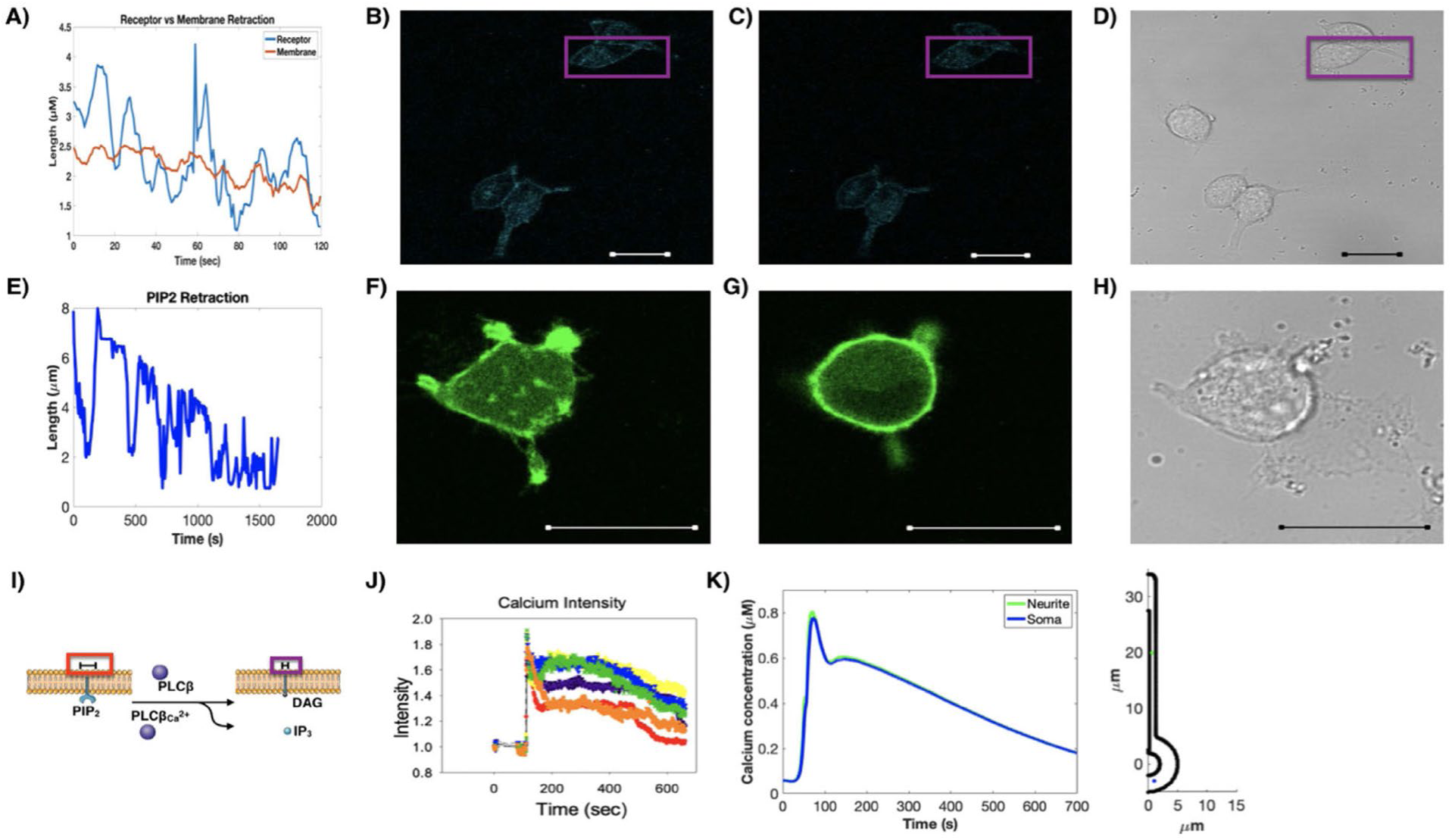
Neurite retraction of PC12 cells upon Gαq stimulation. **A)** Decrease in neurite length of the B2R receptor in PC12 cells (blue) as compared to slow reduction in length of the membrane as followed by phase contrast (orange). **B)** Images of differentiated PC12 cells expressing eCFP-B2R prior to stimulation with bradykinin and (C) 30 minutes after stimulation where purple box shows the retraction data of (A), and the corresponding phase contrast image is shown in (D). **E)** Decrease in neurite length as followed using a fluorescent PI(4,5)P_2_ sensor, PH-PLCδ1 (52) where distinct oscillations are seen. **F)** Images of a differentiated PC12 cells expressing PH-PLCδ1, prior to stimulation, (G) 30 minutes after stimulation with carbachol, and (H) the corresponding phase contrast image are shown. **I)** Cartoon showing the hydrolysis of PI(4,5)P_2_ to Ins(1,4,5)P_3_ and DAG depicting the larger head group of PIP_2,_ denoted by the scale bar (red box), as compared to the smaller DAG that remains in the membrane after hydrolysis, (purple box). This decrease in the size of the membrane bound molecule (PI(4,5)P_2_ to DAG) leads to a local change in tension (denoted by the scale bars). IP_3_ moves into the cytosol and to the ER, causing a release of calcium. **J)** Graph of calcium intensity in neurites of six different cells. This graph shows the lag time after the addition of carbachol, the initial spike in calcium and the slow recovery. **K)** Model replicating the intensity of calcium over time at two spatial locations (in the neurite and in the soma) after stimulation by carbachol. *Inset*: location of traces within the model geometry (in the neurite and in the soma). In all images, the scale bar is 20µm, and n>15 cells.

We calculated the velocity of neurite retraction by analyzing the decrease in length of each neurite at each time point during the experiment (see Methods). These velocities were analyzed for PC12 cells transfected with eCFP-B2R (blue) and for the plasma membrane (orange) as monitored by the phase contrast imaging (Fig. 2A). We find that the velocity of retraction of the plasma membrane is more gradual than that of the receptor suggesting that movement of the receptor towards the soma precedes the membrane.

Activation of the PLCβ pathway through Gαq-GPCRs results in hydrolysis of the signaling lipid (PI(4, 5)P_2_. We followed the change in the level and distribution of (PI(4, 5)P_2_ of the plasma membrane surrounding the soma and neurite during retraction using a fluorescent PI(4,5)P_2_ sensor (i.e. eGFP-PH-PLCδ1). Our data show that PI(4,5)P_2_ moves from the neurite into the soma with a similar retraction velocity as the GPCR (Fig. 2E). However, this movement is oscillatory showing three precipitous drops and recovery during the retraction event. These are three distinct movements of PI(4,5)P_2_during the observed retraction occur at similar times (∼3, 10 and 15 min) for all cells tested, and are interpreted as being due to PI(4,5)P_2_ replenishment during the course of the retraction.

In general, Gαq activation produces an increase in the calcium typified by an initial spike in the first few minutes followed by a slow recovery. We followed the calcium behavior upon stimulation in single cells using a fluorescent calcium indicator, Calcium Green (Fig 1D, F, H and 2J). Correlating calcium responses with neurite retraction shows that retraction occurs even in this initial phase and continues through the duration of the recovery phase.

We constructed a spatial model of the Gαq/PLCβ/PI(4,5)P_2_ signaling pathway using reaction-diffusion equations. The reactions, kinetic parameters, and diffusion constants are given in Tables S1-4 in the Supplementary Material (SOM). The soma of the PC12 cell was modeled as a sphere with a cylindrical neurite as a simplified geometry capable of capturing the key features of these cells. Briefly, the soma was modeled as a sphere with a radius of 5 µm, and the neurite was modeled as a cylinder with a radius of 1.25 µm and length of 30 µm. The endoplasmic reticulum was modeled as a cylinder with a radius of 0.25 µm and length of 25 µm, and the nucleus as a sphere with a radius of 2 µm. Geometric details are given in Table S5. The nucleus was treated as an excluded volume, while the ER was treated as a calcium source. Computational modeling of this pathway in a three-dimensional spatial model using finite elements captured the calcium transients with the same time scale as the experiments (compare Fig. 2J and Fig. 2K). This framework sets the stage for the mechanical coupling of calcium dynamics to neurite retraction.

### Actin remodeling proteins affect the dynamics of neurite retraction

Actin filaments are the major cytoskeletal components of synapses and are the key modulators of neurite plasticity. An actin filament is a dynamic structure in which monomers disassemble from one end and reassemble on the other; a behavior knows as molecular treadmilling. Retraction involves inward movement of the actin structure that defines the neurite shape and so any model of retraction must incorporate actin disassembly. In neurites, actin filament remodeling is known to be associated with the drag forces related to protrusion (26). Additionally, PI(4,5)P_2_ hydrolysis is closely connected to membrane-actin adhesion and membrane tension regulation (22).

Based on our experimental observations (Fig. 2), we coupled PI(4,5)P_2_ hydrolysis and calcium release following carbachol-stimulation with a mechanical model that balances the forces due to actin depolymerization and membrane cortical tension change. Due to the timescale separation between the biochemical signaling dynamics and neurite retraction, we coupled the 3D spatial model of the Gαq/PLCβ/calcium signaling pathway (Fig. lA-B) to an ordinary differential equation model of neurite mechanics, assuming uniform signaling dynamics at extended timescales (Section S1-2 in SOM). Briefly, the mechanical model proposes a force balance acting on the tip of the neurite (34). The force due to the actin cytoskeleton is acting outward, away from the soma and the force exerted by the membrane cortical tension is acting towards the soma. In this case, we assume that the cortical tension is a function of the change in the area of the membrane. This change in membrane area can result from (PI(4, 5)P_2_ hydrolysis (22, 35), from endocytosis of GPCRs (36) and from changes in tension (37). The force exerted by the actin cytoskeleton is modeled phenomenologically as a function of calcium-mediated cofilin and retrograde flow (38). Thus, the net force balance can be written as:

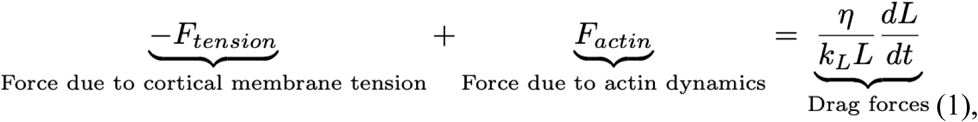

Where

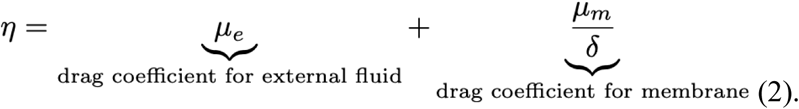

Modeling the neurite as a cylinder with a hemispherical end cap, we obtain the governing equation for the length of the neurite as

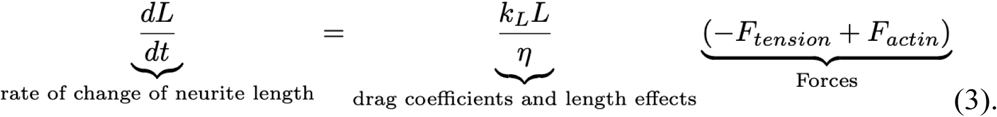

This governing equation combines the force balance above with experimental observations that retraction velocity is proportional to neurite length. The force due to membrane cortical tension and actin dynamics depend on equations for and F_actin,_ respectively, given as

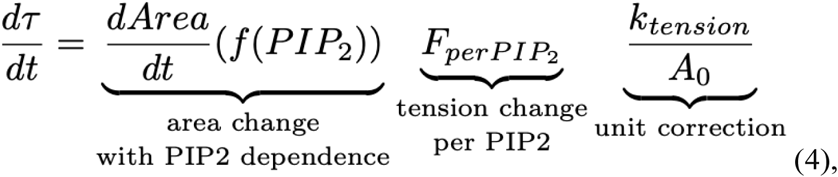

and

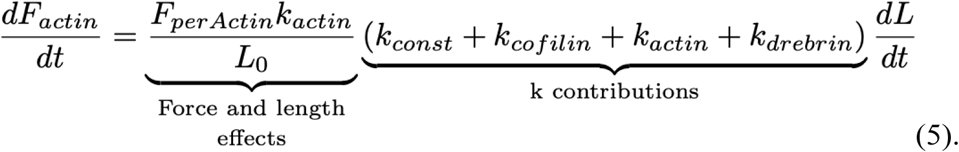

This is a phenomenological model with a combination of parameters from literature and constants fit to the experimental results, (see SOM). In particular, Eqs. (4) and (5) have dependence on f(PIP2) and g(CofilinAct), which are sigmoidal functions that trigger in response to (PI(4, 5)P2 and activated cofilin, respectively(39).

Using this model, we simulated calcium-mediated neurite retraction in control cells. We found that the model is able to capture the dynamics of neurite retractions events (Fig. 3C). These models utilized experimental retraction data collected using differentiated PC12 cells transfected with low amounts of mCherry-actin to fit the phenomenological constants (Fig. 3). Effectively, the model predicts that when the force exerted by actin is matched by the force due to tension, there is no change in the length of the neurite. Upon increase in PI(4,5)P_2_ hydrolysis and calcium release into the cytosol, the forces due to membrane cortical tension increase while the forces exerted by the actin decrease and the neurite pulls back.

**Figure 3:**
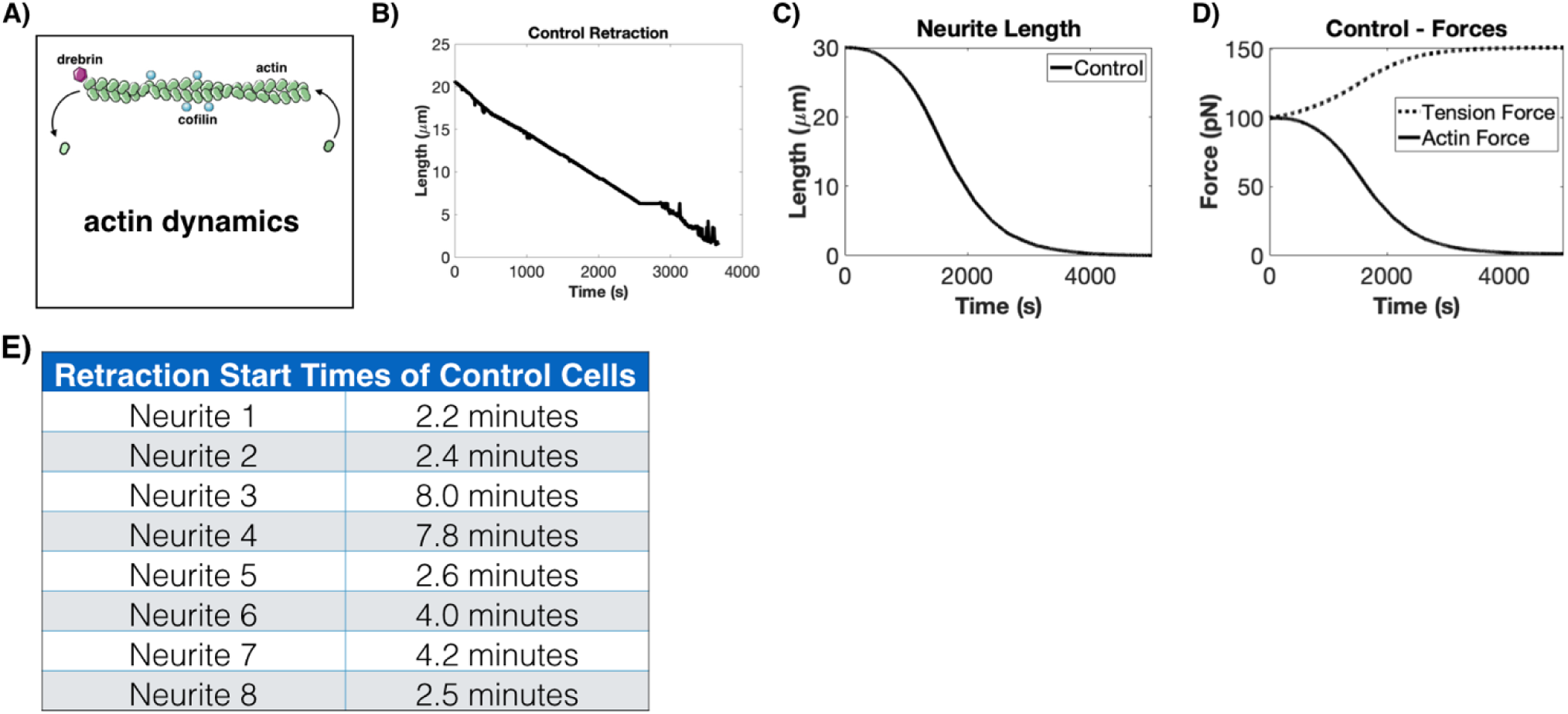
Modeling of neurite retraction. **A)** Simplified schematic of actin dynamics depicting assembly/disassembly as a combination of effects due to actin, cofilin, and drebrin concentration**. B)** Example of the retraction in response to carbachol stimulation of differentiated PC12 cells transfected with mCherry-actin. Identical behavior was seen over an average of 8 neurites shown in the table **E). C)** Computational results for neurite retraction determined from the described model. **D)** Calculation of force plots due to increased tension in response to PI(4,5)P_2_ hydrolysis, internalization of ligand-bound GPCRs, and tension effects. The force due to actin decreases and switches direction in response to actin breakdown and retrograde flow. **E)** Table showing the variation of retraction times within the same condition.

If this prediction is true, then we should be able to alter neurite retraction behavior by altering the expression of actin-related proteins or by membrane tension separately. We next probed neurite retraction dynamics by varying the concentration of select actin-related proteins such as cofilin, drebrin, and actin itself both in experiments and modeling.

### Actin modulators regulate neurite retraction velocity

We followed actin disassembly in real time during carbachol-stimulated retraction using mCherry-actin in control cells and cells over-expressing cofilin or drebrin (Fig. 4). Cofilin increases severing of actin filaments increasing actin filament breakdown while drebrin stabilizes actin filament inhibiting disassembly (40–43). Fig. 4 D-L are screen shots of the time lapse video made during retraction where Fig 4. D-F are representative cells prior to stimulation with carbachol, Fig. 4G-I show the same cells at the beginning of retraction, and Fig. 4J-L show the cells at longer times. Movies of retraction (*SOM2*) were analyzed to obtain the retraction velocities during the three different conditions (Fig. 4M-P). This quantification enables the comparison of neurite retraction dynamics of actin in our control versus actin when cofilin and drebrin are overexpressed. As expected, overexpression of cofilin enhances the rate of neurite retraction (Fig. 4M versus Fig. 4N), and complete retraction results in cells with a trapezoidal morphology rather than the circular one typified of undifferentiated PC12 cells. This altered morphology likely stems from the lower number of actin filaments throughout the cell due to cofilin overexpression (44).

**Figure 4:**
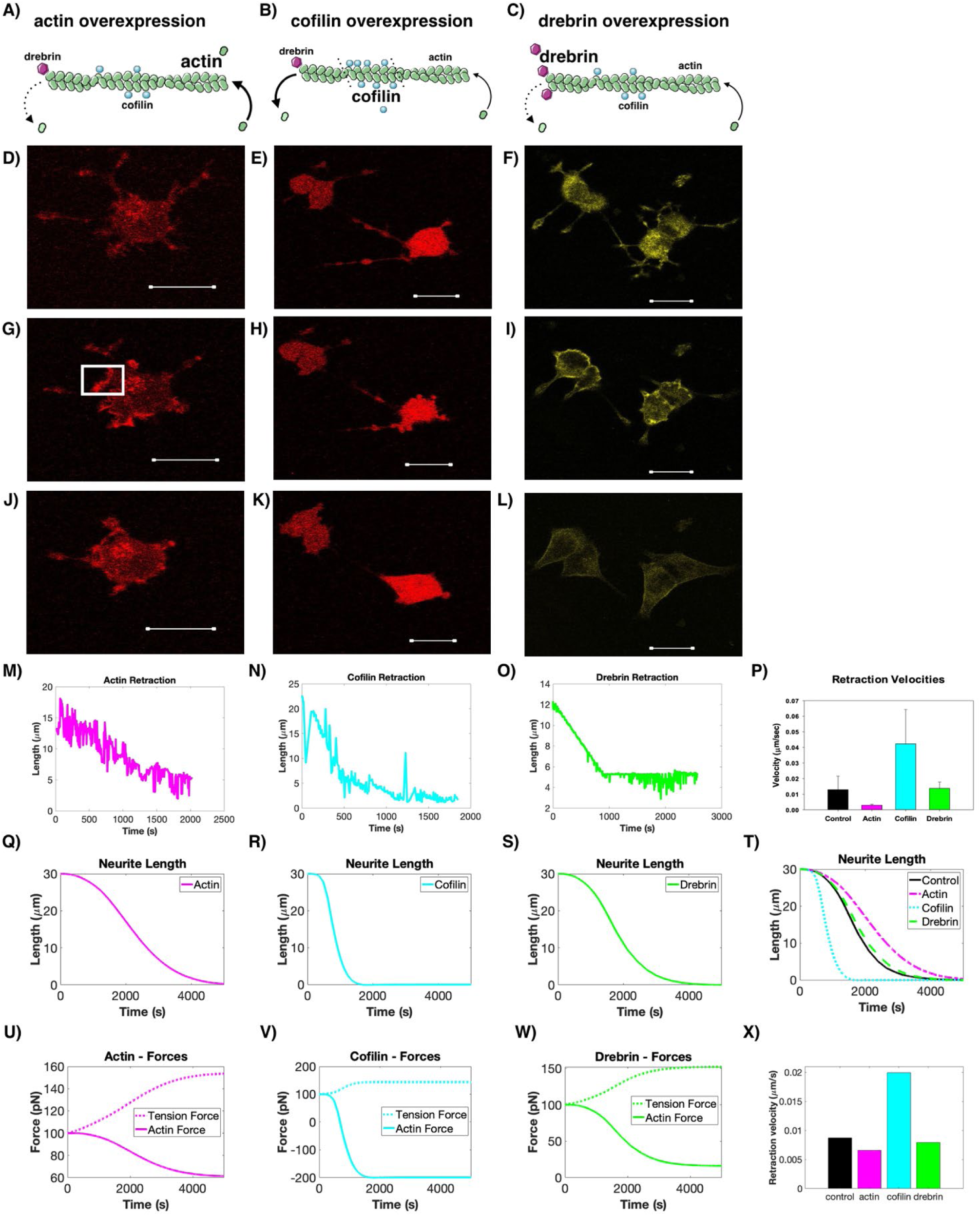
Experimental validation of model predictions. **A-C)** Schematic of the experimental study to test the actin force model where mCherry-actin, mCherry-cofilin or eYFP-Drebrin was expressed at varying levels in PC12 cells using increasing amounts of DNA in the transfections. The corresponding images shown directly below. While cofilin activity is mediated by calcium, actin and drebrin dynamics depend on local concentrations. *Column 1* (panels A, D, G, J, M, Q, U) are results from studies following mCherry-actin expressed in PC12 cells. The increase in actin, as estimated by western blotting, reduces the change in the actin force. Panel **D** is an example image of cells before carbachol stimulation. **G** is 15 minutes post-stimulation showing the beginning stage of actin breakdown indicated by the white box, and **J** is 30 minutes post-stimulation. A representative retraction velocity curve is shown in **M**. Panels **Q** and **U** are the results of modeling the retraction velocity (Q), due to a tension force and actin force (U). We note that the overexpression of actin leads to less of a decrease in actin force since the increased actin concentration slows overall actin breakdown. *Column 2* (panels B, E, H, K, N, R, V) shows how cofilin overexpression creates a large negative actin force to increase the rate of disassembly as shown in the schematic (B). Panel **E** is an example image of cells expressing cofilin-RFP before carbachol stimulation, **H** is 7 minutes post-stimulation, and **K** is 16 minutes post-stimulation. A representative retraction velocity curve is shown in **N**. Panel **R** and **V** are the results of modeling the retraction velocity (R), due to the tension force and a larger negative actin force (V), since increased cofilin concentration increases actin breakdown. *Column 3* (panels C, F, I, L, O, S, W) shows how drebrin overexpression slows the change in actin force as shown in the schematic (C). Panel **F** is an example image of cells expressing drebrin-eYFP before carbachol stimulation. **I** is 15 minutes post-stimulation, and **L** is 30 minutes post-stimulation. A representative retraction velocity curve is shown in **O**. A representative retraction velocity curve is shown in **O**. Panels **S** and **W** are the results of modeling the retraction velocity (S), due to the tension force and actin force (W). We note that increased drebrin concentration resists actin breakdown. Panels **P**, **T** and **X** are compiled retraction velocities obtained experimentally (P), control (n=4), actin(n=4), cofilin(n=3), drebrin(n=4) showing standard error, and computationally (T and X), showing the very close correlation between experimental and theoretical results. All scale bars = 20 µm.

We expected that transfection of the cells with drebrin would strongly impede neurite retraction in the presence of carbachol because drebrin is known to bind assembled actin filaments (F-actin) and stabilize them (41, 45, 46). Low amounts of actin allowed a sustained linear decrease (Fig. 4O) whose compiled average is within error of mCherry-actin along (Fig. 4P). These data support the idea that disruption of the intact actin network and its dynamic modeling when impeded by cofilin overexpression (increased severing) or by drebrin overexpression (inhibition of disassembly and remodeling) are key driving forces in governing retraction dynamics.

We modeled overexpression of cofilin, drebrin, and actin in the mechanical framework by altering the F_actin_ terms (Eq. 5) to represent the known effects of these proteins through the ‘k’ contributions for each protein. We modeled actin force overall as a stress-strain relationship, where the ‘k’ terms captured the contribution of each protein to overall actin force dynamics. For example, cofilin expression was incorporated in our constitutive equation as an increase in the total cofilin available for calcium activation, which therefore causes a faster decrease in F_actin_ by increasing the magnitude of the rate of change of F_actin_ through the k_cofilin_ term. As a result, our model predicts that neurite retraction velocity is faster for cofilin overexpression compared to control (Fig. 4R,V). Similarly, we modeled drebrin overexpression by increasing drebrin concentration within the k_drebrin_ term, consistent with experimental observations that a higher drebrin concentration slows actin depolymerization and thus slows the decrease in F_actin_ (41, 46). Therefore, our model predicts that drebrin overexpression slows neurite retraction velocity compared to control (Fig. 4S,W). Actin overexpression can be modeled in a similar fashion as drebrin and is captured through the k_actin_ term. Thus, increasing actin concentration is essentially increasing the actin-to-cofilin ratio and therefore slowing the decrease in F_actin_. Our model predicts that actin overexpression will slow neurite retraction velocities compared to control (Fig. 4Q,U).

The results in Fig. 4 show that by varying cofilin, drebrin, and actin concentrations, we predict different actin force dynamics that translate to different neurite retraction rates. Specifically, we predict that cofilin, actin, and drebrin overexpression leads to more rapid, slower, and slightly slower neurite retraction velocities compared to control, respectively. Our modeling results agree with the experimental observations and suggest that actin disassembly and inhibition of reassembly are necessary for retraction. To support these studies, we quantified experiments that measured retraction of control cells and cells that overexpress actin, cofilin and drebrin. The experimental velocities of the retractions are plotted in Fig. 4P and validate our model predictions in Fig. 4X.

### Neurite retraction can be driven by membrane tension

Given that there are two contributions to the force acting on the neurite, one from actin and the other from membrane cortical tension, we next asked whether neurite retraction could be driven by changes to the membrane cortical tension alone. We directly tested this idea using hyper-osmotic stress to increase membrane tension. Increasing the hyperosmotic stress from 300 to 600 mOsm on cultured PC12 cells resulted in the same retraction behavior and timescale as seen with carbachol stimulation (Fig. 5A). To determine the change in membrane tension induced by this change in osmolarity, wemeasured tension by atomic force microscopy (AFM). This method allows us to assess the stiffness of cells by measuring the amount of deflection experienced by a cantilever with known spring constant as it indents the surface of the cell. For undifferentiated, differentiated, and retracted PC12 cells, AFM measurements were taken at the base of the neurites or retracted neurites. The stiffness of each cell was determined by averaging the stiffness values at three separate locations on each cell in order to combat the inherent heterogeneity in cell structure while ensuring no localized damage from a previous measurement. Undifferentiated cells were the stiffest condition tested, followed by retracted and then differentiated (Fig. 5B).

**Figure 5:**
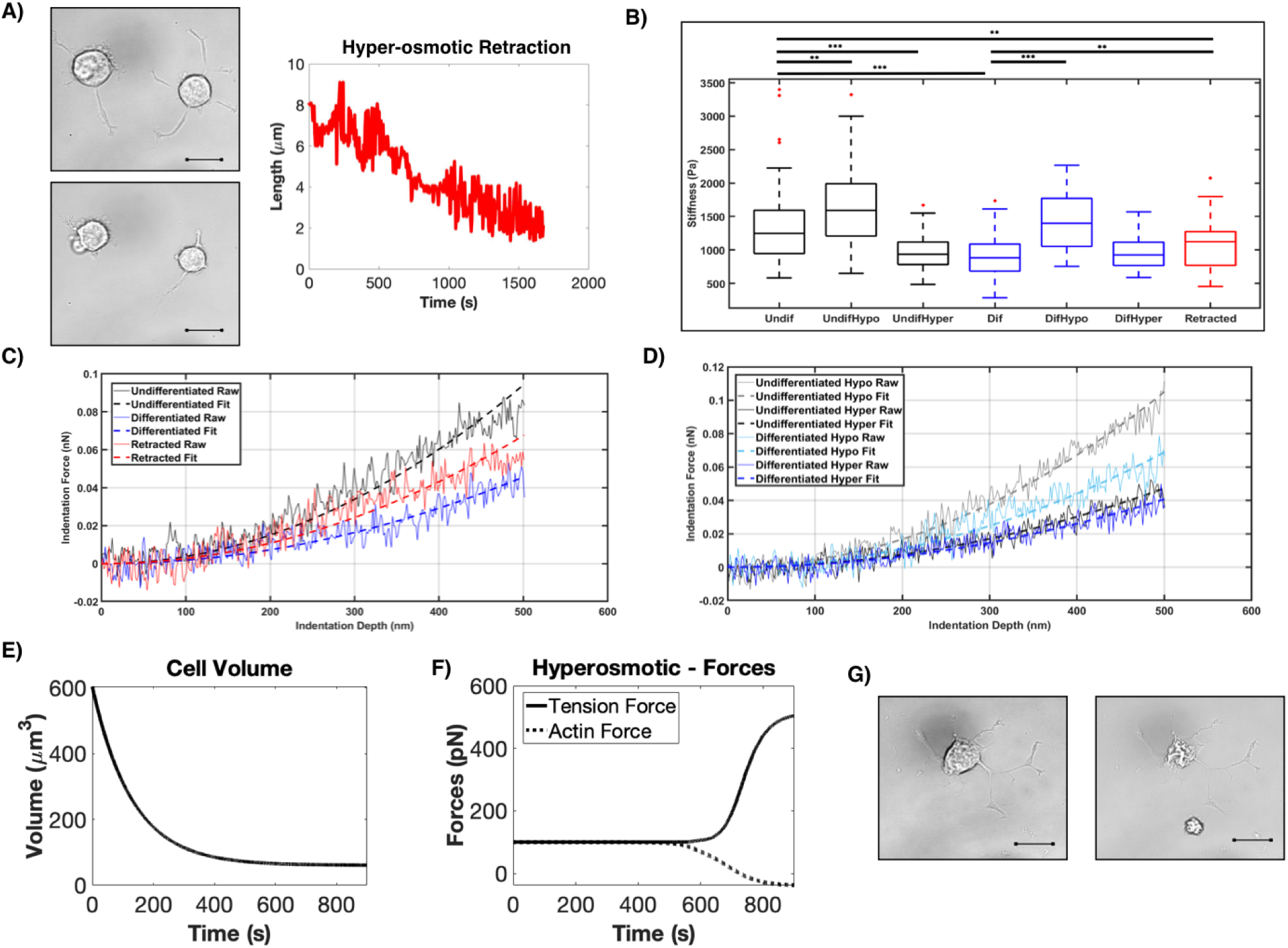
Neurite retraction can be induced by membrane tension in response to hyper-osmotic stress. **A)** Phase contrast images of PC12 cells under isotonic conditions (300 mOsm, top left) and hyper-osmotic conditions (450 mOsm, bottom left) after 10 minutes). The corresponding neurite retraction curve for the cell on the left is shown to the right. **B)** Cell stiffness as measured by atomic force microscopy of undifferentiated and differentiated PC12 cells under isotonic conditions and cells subjected to hypo-osmotic (150 mOsm) or hyper-osmotic (450 mOsm) stress where the number of cells n =14 – 46, and were * = p < 0.05, ** = p < 0.01, ***= p < 0.001. **C-D)** Force curves representative of the average stiffness values in B with automated Hertz model fitting (see methods) over 500nm indentation for untreated cells (left) and hypo-osmotic/hyper-osmotic treated cells (right). **E-F)** Hyperosmotic stress is modeled as an instantaneous increase in membrane tension due to the reduction in volume-to-surface area ratio, which causes a membrane tension increase and a smaller cytoskeletal force (E), and triggers neurite retraction (F) in the absence of ligand input. **G)** Phase contrast images of a differentiated PC12 cells before the application of osmotic stress (left) and after increasing the osmolarity from 300 to 600 mOsm in calcium-free media. Note the loss in cell volume seen by membrane puckering but without retraction (right)

We also used AFM to determine the stiffness of cells subjected to hypo-osmotic and hyper-osmotic stress for undifferenced and differentiated PC12 cells. Hypo-osmotic and hyper-osmotic stress, which leads to cell swelling and cell dehydration respectively, results in an increase and decrease in cell stiffness, respectively (Fig. 5B). Force curves that are representative of the average stiffness value for each condition show a smooth Hertz model relationship between indentation depth and applied force as expected (Fig. 5C-D). Taken together, these studies show a direct correlation between cell stiffness and osmotic stress, which is also correlated to retraction.

Within our model, we can also probe neurite retraction dynamics by varying the contribution of tension, the component, of the mechanical model. Membrane cortical tension is altered under hyperosmotic conditions when water expulsion from the cell leads to a reduction in volume-to-surface area ratio. Therefore, to model hyperosmotic conditions instead of a carbachol stimulus, we predict that this volume-to-surface area ratio change causes an increase in membrane cortical tension. Therefore, we model a slightly elevated membrane cortical tension initial condition that triggers a related force (Fig. 5E) and neurite retraction (Fig 5F). We also see a change in actin force, triggered due to the coupling between actin and tension in the model, which is also predicted experimentally due to the necessary actin reorganization during retraction caused by hyperosmotic conditions. Therefore, our model predicts neurite retraction in the absence of a ligand stimulus with hyperosmotic conditions (Fig. 5F), indicating that membrane cortical tension is a key component governing neurite retractiondynamics.

We tested whether the retraction that results from osmotic stress is convergent with the Gαq/PLCβ pathway. Keeping in mind that membrane compression regulates calcium flux to drive actin dynamics in model systems (47–49), we determined whether hyper-osmotic stress will open calcium channels to allow divalent cation influx to relieve the osmotic stress. Using a fluorescence calcium indicator, we found that increasing the osmotic strength from 300 to 600 mOsm results in a small increase in intracellular calcium that is ∼10% of the increase seen for ACh stimulation. To verify that calcium is mediating retraction, we removed extracellular calcium from the media and found that even though increasing the osmotic strength through the addition of KCl caused membrane folds and puckering, consistent with a loss in cell volume, neurite length remained constant (Fig. 5G). This result indicates that the two types of retraction pathways converge on calcium levels, those initiated by Gαq/PLCβ stimulation and those initiated by extracellular calcium influx by tension sensitive calcium channels.

### Prolonged exposure to Gαq agonists leads to nerve ring disruption in *C. elegans*

To determine whether we could disrupt neuronal connections and retract neurites in a neuronal network, we used the optically clear model system, C. *elegans.* We first used a strain of *C. elegans* (QWl166) that expresses an integrated fluorescent calcium sensor G-CaMP throughout the nervous system. This system allows us to visualize by fluorescence, the neurons that show increased calcium levels in response to acetylcholine stimulation. This specific strain has fluorescence in every neuron in the nerve ring allowing us to look at the ring as a whole, as well as fluorescence in the entire ventral nerve of the worm that extends the entire length of the worm. We focused on acetylcholine signals emanating from neurons within the nerve ring located in the head of the worm and along the ventral nerve cord (Fig 6A). Normal signal transmission can be seen in the images in Fig 6B. For these studies, we only wished to determine whether there is broad neuronal rupture and retraction within the nerve ring and along the ventral nerve which would be indicated by the lack of fluorescence in the center in the ring and dark spots along the ventral nerve. In every worm tested (n=10), we find neuronal rupture with acetylcholine stimulation that occurs in the nerve ring close to the organism’s mouth (Fig. 6C). Additionally, we see a distinct pilling of the ventral nerve that displays thinning and clustering, and portions that are no longer fluorescent indicting large-scale neuronal damage.

**Figure 6:**
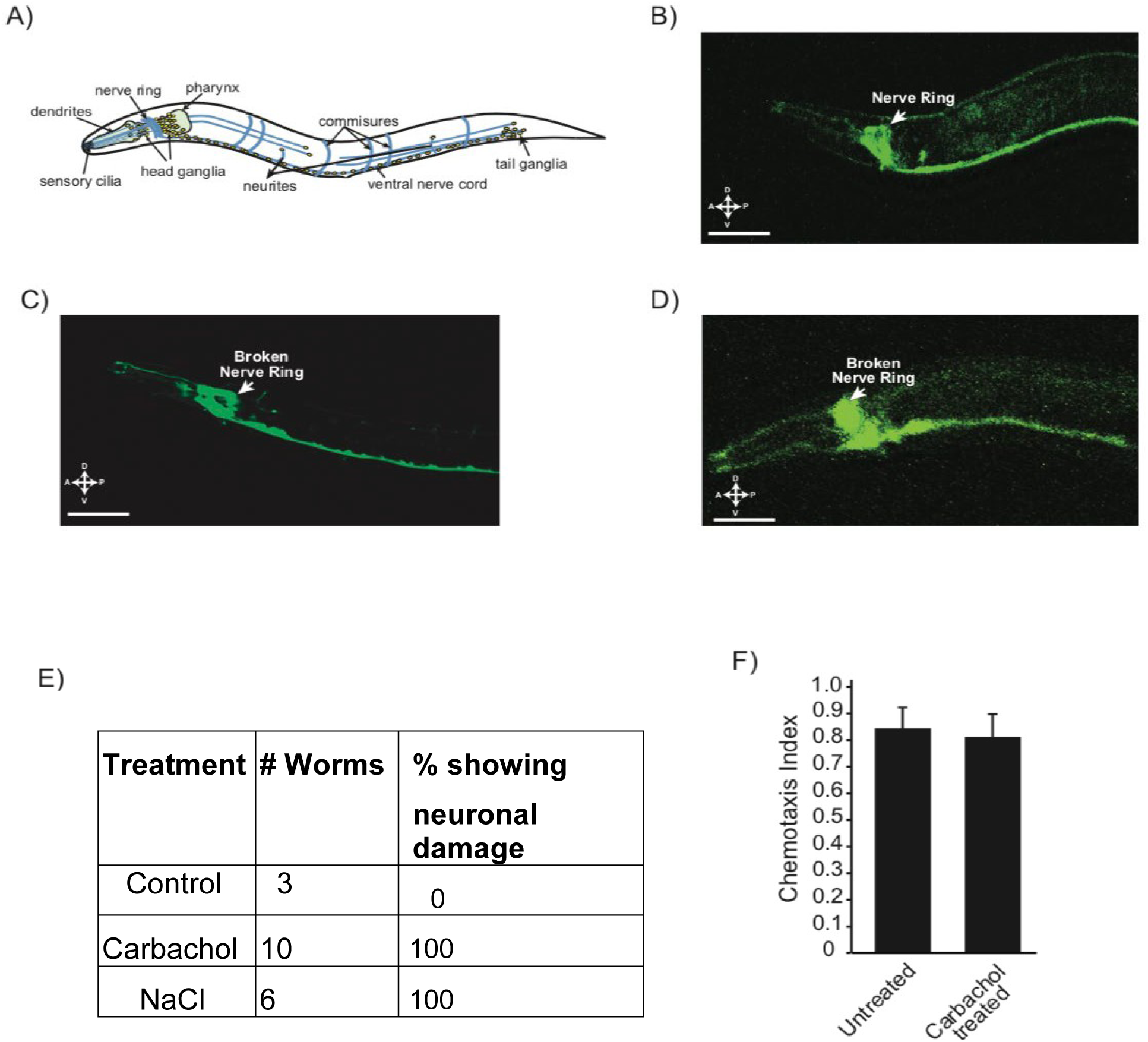
Neurite retraction induced by membrane tension is seen in the neural network of ***C.elegans***. **A)** General schematic of the nervous system of C. *elegans.* **B**) Untreated C. *elegans* that expresses GFP G-CAMP through the nerve ring and ventral nerve. **C**) A representative worm with 1mM carbachol for 30 minutes showing a broken nerve ring and pilling ventral nerve. **D**) A representative worm subjected to hyper-osmotic stress showing a broken nerve ring. **E**) Table showing the compiled results of each experimental condition and the number of resulting nerve rings that were broken. **F**) Results of a chemotaxis assay testing movement of C. *elegans* with and without lmM carbachol treatment.

We confirmed the connection between neuronal network disruption produced by extended acetylcholine stimulation to membrane tension by subjecting the worms hyper-osmotic stress (increasing the salt solution concentration by 50%). After osmotic stress, these organisms showed a lack of fluorescence in the nerve ring indicating rupture and retraction of neurons. Also, we see much larger segments of the ventral nerve that have been disrupted with less pilling than that seen for acetylcholine stimulation (Fig. 6D). The somewhat different morphological effects on the ventral nerve seen in osmotic stress is most likely due to the ability of the stress to be distributed uniformly along the length of organism as opposed to carbachol whose receptors are concentrated mostly in the mouth.

To support neuronal disruption of the worms by acetylcholine, we followed changes in movement of the whole organisms using a chemotaxis assay with and without acetylcholine stimulation. These studies were also done using the *C.elegans* QWl166 strain. We see that after treatment with acetylcholine 83% of the worms moved towards the attractant while the remaining 17% did not move at all (Fig. 6F).

## DISCUSSION

In order for an organism to learn, neurons must break and reform connections with neighboring neurons and disruptions in these processes underlie learning-based neurological diseases and neurodegeneration. Despite the importance of synapse severing and formation, the mechanisms that underlie neural plasticity are not well understood (3). Previous studies of neural plasticity have mainly focused on contributors to neurogenesis in hippocampal neurons and cultured drosophila neurons, such as neuronal stem cells (50). Here, we have defined the synaptic breakage and neurite rupture in neural plasticity in terms of membrane tension and calcium-induced retraction that will allow for predictable actin remodeling.

Calcium is a crucial component for the growth, differentiation, and survival of neurons (12). Calcium helps to regulate the differentiation of specific neuronal types as well as their migration through the body, and calcium levels are directly liked to neurodegenerative diseases such as Alzheimer’ s (50). We followed neuronal cell response to extended calcium signals that may mimic dysfunctional states. Unexpectedly, we find that extended calcium levels resulted in complete neurite retraction, and this retraction can be seen by following the plasma membrane, by a GPCR coupled to Gαq, and by PI(4,5)P_2_. These data show that the Gαq signaling pathways are intimately involved with the mechanical properties of the cell.

The connection between calcium signals from GPCR/Gαq and cell mechanics was supported using fluorescent-tagged actin and by measuring retraction when overexpressing cofilin, an actin depolymerizing protein, or drebrin, an actin monomer recruitment protein. Overexpressing cofilin at a level 6x higher than endogenous levels allows for much faster velocities of neurite retraction initiated by GPCR-Gαq/Ca^2+^ after an initial lag period. These observations correlate well with cofilin’s ability to depolymerize actin, but shows that activation of the Gαq/PI(4,5)P_2_/Ca^2+^ pathway is a necessary prerequisite. In contrast, cells overexpressing drebrin show a similar retraction rate as endogenous which correlates to its ability to stabilize actin monomers (41). This observation indicates that the behavior we are viewing is solely due to actin depolymerization without contributions of the polymerization events. Thus, while Ca^2+^ signals drive retraction, actin plays a passive role.

The coupling between calcium signaling and mechanical forces has been observed in cell motility and in dendritic spines during long-term potentiation, astrocyte calcium signaling, and during neurite protrusion and formation (51). Based on these studies, we postulated that neurite retraction results from changes in cortical tension brought about by receptor endocytosis, (PI(4, 5)P_2_ hydrolysis and IP_3_ generation, and an increase in intracellular calcium, and formulated a mechanochemical pathway to test this idea. We found that indeed, a force balance between cortical tension and actin-mediated forces, both of which are mediated by intracellular calcium levels, is sufficient to capture the dynamics of neurite retraction. This idea was explicitly tested by measuring changes in membrane stiffness with osmotic strength and by altering the effects of different actin remodeling proteins. We also found that the effects of actin and cortical tension could be uncoupled to a certain extent, pointing towards multiple mechanical pathways that could exquisitely regulate neurite retraction. We acknowledge that the proposed model is phenomenological, and while it captures the key physics of the processes underlying neurite retraction, future studies will focus on elaborating on some of the phenomenological relationships proposed here.

Our results show that acetylcholine-induced calcium signals may have a dramatic impact on neuronal morphology and synaptic connections, and this idea was demonstrated in *C. elegans.* Neurite formation and extension is essential to transition from neuroblast to mature neuron and *elegans* are an excellent model system to understand neurite formation and retraction (Carr et al 2016). The site of nascent neurite outgrowth is determined by cell intrinsic factors that orchestrate the localized regulation of actin and microtubules (van Beuningen SFB, 2015, Flynn KC, 2012) and in developing neurons, these factors are polarized at sites of neurite outgrowth in response to extracellular cues (Adler CE, 2006, Randlett O 2011). Our finding that the neural network of this organism can be modified by Gαq activation or increased mechanical tension, may form a basis for studies that better describe neural rewiring paradigms.

In summary, our studies indicate the role of acetylcholine in facilitating neurite retraction. Further research on how this pathway interacts with other pathways to coordinate remodeling with such spatio-temporal precision will provide insights into understanding the molecular basis of neurodegeneration.

## Supporting information

Data associated with modeling and experimental data

## ACKNOWLEDGEMENTS

This work was supported by NIH GM116187 (SS) and AFOSR MURI FA9550-18-1-0051 (PR). We thank various members of the Rangamani and Scarlata labs for valuable discussion and manuscript review. We would like to thank the UCSD Interfaces Graduate Training Program and the San Diego Fellowship for funding and support for MB. MB was also supported by AFOSR MURI grant number FA9550-18-1-0051 to PR and a grant from the National Institutes of Health, USA (NIH Grant T32EB009380).

## AUTHOR CONTRIBUTIONS

KP and MB carried out all of the experimental and theoretical studies and assisted in writing. WL and QW assisted in some of the experiments. JS help to design, oversee some of the studies and writing. PR and SS were responsible for conceptualizing and designing the experiments and writing the manuscript.

## MATERIALS AND METHODS

### Cell Culture, transfection and differentiation

PC12 cells, which are derived from rat pheochromocytoma (ATCC CRL-1721), were cultured in 35mm or 100mm poly-d-lysine coated petri dishes using Dulbecco’s Modified Eagle’s Medium (DMEM) (Gibco) with 10% heat-inactivated horse serum (Gibco), 5% fetal bovine serum (Atlanta Biologicals), and **1**% penicillin/streptomycin. The dishes were incubated with 5% CO_2_ at 37°C. Cells were transfected using different amounts of plasmid based on the concentration tested using a NanoDrop. Cells were transfected using Lipofectamine 3000 (Invitrogen) following the protocol of the manufacturer. The media used in transfection was the same DMEM culture media with the exception of antibiotics, to increase transfection efficiency and the media was changed back to normal culture media after 24 hours. Cells were differentiated using media that contained DMEM, 1% heat-inactivated horse serum and 1% penicillin/streptomycin. Added to this would be a 1 to 1000 ratio of l00ng/µl nerve growth factor (NGF) (Novoprotein). This media is added to the cells for at least 48 hours and up to 96 hours to achieve long neurites.

### Plasmids and Maxi/Mini prep

Fluorescent-tagged plasmids were obtained and maxi/min prepped using Qiagen kit and following the manufactures guidelines. The plasmids were obtained from Addgene. Actin #54967 derived by Michael Davidson at Harvard Medical School, Drebrin #40359 derived by Phillip Gordon-Weeks at King’s College of London, Cofilin #51279 derived by James Bamburg at Colorado State University, Pleckstrin homology (PH)-domain of PLC8 # 21179 derived by Tobias Meyer at Stanford University, and the bradykinin type 2 receptor was modified from the construct provided by Porf. Leed-Lundber, Univ Texas San Antonio.

### Calcium Green Staining

Prior to imaging, cells are washed with HBSS and then a 1/200 mixture of HBSS and Calcium Green (Thermofisher) is added and allowed to incubate for **1** hour before imaging.

### Preparation of Carbachol

Carbachol in powdered form was obtained from Sigma Aldrich. It was dissolved in water to a final concentration of l mM at a volume of l0 mL and aliquoted into 500µL portions to be used for each experiment. The solutions were kept at −20°C.

### Fluorescent Microscopy/ Stimulation of Cells

Fluorescent imaging was done using a Zeiss LSM510 inverted confocal microscope. Imaging was carried out at least 48 hours post transfection and differentiation. The cells were grown and imaged in a MAT-Tek 8 chamber glass bottom plates or a MAT-Tek 35mm glass bottom dish. Once a single cell was found visually that expressed the plasmid or stain, the microscope would be switched to the correct wavelength and laser intensity. A time series image was then started and stopped after 10 frames, the cells would then be manually stimulated by adding carbachol to the dish to achieve the desired final molarity, and the time series video was then started immediately after adding the Carbachol.

### Osmotic Stress and EDTA studies

Osmotic stress studies were done using calcium free isotonic media for imaging with the addition of KCl at different concentrations to achieve osmotic stress. Studies that include EDTA, (J.T. Baker) cells were osmotically stressed and EDTA was added to the dish to give a final concentration of 0.5 for partial neurite retraction and 1μM for full neurite retraction.

### Plasmid-specific studies

Calcium green and enhanced green fluorescent protein (eGFP) studies were imaged using an argon laser at 488nm. Red fluorescent protein (RFP) and mCherry were imaged using argon ion and HeNe lasers at 543nm. YFP and CFP were imaged using argon ion and HeNe lasers at a wavelength of 545nm. Multi-track imaging can also be done combining each of these set ups. Images were taken by alternating between probes when data for more than one are being monitored, and these images are subsequently merged.

### Atomic Force Microscopy Stiffness Measurements

Live cells were probed utilizing an MFP-3D-BIO atomic force microscope (Asylum Research) and a DNP cantilever (Bruker) with nominal spring constant 0.06 N/m[l]. The cantilever was calibrated before each measurement to ensure accuracy. Cells were seeded on 60 mm poly-d-lysine coated petri dishes using Dulbecco’s Modified Eagle’s Medium (DMEM) (Gibco) with 10% heat-inactivated horse serum (Gibco), 5% fetal bovine serum (Atlanta Biologicals), and 1% penicillin/streptomycin. After 24 hours of recovery cells were differentiated using media containing DMEM, 1% heat-inactivated horse serum and 1% penicillin/streptomycin. Added to this would be a 1 to 1000 ratio of lO0ng/µl nerve growth factor (NGF) (Novoprotein). This would be added to the cells a minimum of 24 hours before measurements are taken.

Cells were viewed two days after plating and one day after treatment with nerve growth factor (NGF). Cells with minimal cell-cell contact were selected to reduce the mechanical impacts of cell communication. Three force curves in separate perinuclear regions with cantilever velocity of 2µm/s and trigger point lnN were taken for each selected cell. Measurements were taken within 30 minutes of removal from the incubator to facilitate cell viability.

The stiffness of the measured cells ***E*** was determined from the force curve data utilizing the Hertz model for conical cantilever tip geometry:

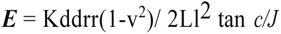

where ***k*** is the cantilever spring constant, dis the cantilever deflection, vis the poison’s ratio (0.5 used for an assumed incompressible material), ***A*** is the sample indentation depth, and ***<P*** is the half-angle of the conical cantilever tip (35°). The force curves were processed using a custom MATLAB code that fits the indentation curve over a 500nm range after manual selection of the initial contact point.

### Fluorescent Imaging of C. elegans

Worms were transferred into a microcentrifuge tube containing l mM carbachol and were submerged in the tube for 30 minutes at room temperature or 100mM NaCl solution for 10 minutes. After 30 or 10 minutes respectively, the worms were removed from solution and placed on a glass slide with a thin layer of agar on it for the worms to lay on. Excess carbachol was wicked up from the pad and the worms were paralyzed using 25M levamisole that does not a affect our pathway of interest. The worms were then imaged on an inverted confocal microscope and were visually checked for neurite rupture and retraction.

### Chemotaxis Assay of C. elegans

Worms were transferred to a new seeded plate 24 hours prior to the experiment. The day of the experiment an unseeded plate is marked with a dot in the center and two circles on either ends of the plate with the “C” for control of DI water and “D” for diacetyl which is the attractant. Using approximately 1mL of s. basel the worms are washed into a microcentrifuge tube and allowed to pellet for 5-10 minutes. Once pelleted, the worms are washed with s. basel two more times and finally with water. 10μL of the suspended worms are transferred to the experimental plate previously made and the excess water is removed by wicking. The plate is then incubated for 45 minutes at 20°C, and after, the number of worms that have moved either to the attracted or control is counted.

## Model development - Computational Model

### S1. Model Assumptions

The mechanochemical model developed here is based on the following assumptions:

1. The reaction in the biochemical model were based on [17693463], [17483174], and [10.1101/161950]. These are models of calcium signaling in cardiomyocytes and vascular smooth muscle cells. We assume that the signaling dynamics of the/PIP_2_/IP_3_ pathway are similar enough to PC12 cells to use these reactions and parameters in our model.
2. We assume an axisymmetric 2D geometry, which imposes rotational symmetry. We assume this idealized geometry as a simplification of the PC12 cell to focus on the underlying mechanochemical effects.
3. Geometry specification - We model a single neurite of 30 m based on experimentally observed neurite lengths. We include the ER along the neurite and include the nucleus as excluded volume. Additional experimental imaging can provide more information on the ultrastructure within these PC12 cells.
4. Mechanical model framework - We developed a mechanical model that has two main factors, tension and actin, based on experimental observations of hyperosmotic stress tests and actin-related protein overexpression experiments. We used a phenomenological model for connecting signaling and mechanics due to the complexity of the underlying mechanisms and the lack of information on a more complicated signaling framework. We model the mechanical portion as a system of ODEs coupled to the signaling model.

### S2. Biochemical signaling network in neurites

We developed a 2D axisymmetric spatial model of an idealized PC12 cell with a single neurite. We modelled the biochemical signaling pathway in response to G protein coupled receptor (GPCR) activation due to Carbachol stimulation, specifically involving PIP_2_ hydrolysis, IP_3_ production, and Ca^2+^ release from the endoplasmic reticulum. The specific network can be seen in Fig 1A-B.

#### S2.1 Reaction-diffusion dynamics

Signaling dynamics for each species is given by reaction-diffusion dynamics in both the volumes and on membranes. Reaction-diffusion dynamics of a species Sx, xɛ {1,…,18} are given by

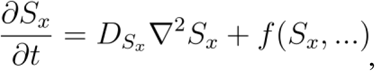

where f(S_x_,…) are any reactions regarding species S_x_. For volumetric species, flux into or out of the volume is given by a membrane flux boundary condition written as

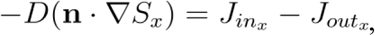

where J_in_ and J_out_ are the various flux reaction into and out of the volume, respectively.

#### S2.2 Note on length scale factors

When transitioning between a volume and membrane, there is a characteristic length scale captured by the volume-to-surface area ratio, ‘n.’ This length scale is used to convert between a volumetric reaction to a membrane boundary flux.

#### S2.3 Signaling species

We have three separate volumetric compartments (extracellular space, cytoplasm, and endoplasmic reticulum), and two boundaries (plasma membrane (PM) and endoplasmic reticulum membrane). We give a general overview of the various signaling species in each volume and on each membrane. Table S1 lists all of the signaling species and initial conditions.

#### S2.4 Extracellular space (ECS)

We model carbachol stimulation of PC12 cells, so we include carbachol as a ligand (L) in the extracellular space. We model the extracellular space as a large region around the PC12 cell.

#### S2.5 Plasma Membrane (PM)

Carbachol (L) binds to G protein coupled receptors (Receptor) on the plasma membrane, forming a bound complex (LR). G proteins are bound to GDP (GD) or GTP (GT). GPCRs can bind GDP (R_GD) and when activated by the ligand form L_R_GD. This activated complex then can become phosphorylated (L_R_GD_P) and is eventually endocytosed and recycled. We do not explicitly model this endocytosis pathway. GTP bound G protein then activates (PLC) to form PLC_GT. also binds Ca^2+^ (PLC_Ca) and can be bound by both Ca^2+^ and activated G protein (PLC_Ca_GT). The two forms of activated catalyzes the hydrolysis of PIP2, forming IP_3_ in the cytoplasm and leaving DAG in the membrane.

We also assume that there are PMCA pumps along the plasma membrane that work to pump calcium out of the cytoplasm and into the ECS.

#### S2.6 Cytoplasm (Cyto)

When PIP_2_ is hydrolyzed, IP_3_ is released into the cytoplasm where it can be degraded into IP_3_-deg or moves to the endoplasmic reticulum membrane to trigger IP_3_R activation. IP_3_Rs release Ca^2+^ from ER stores into the cytoplasm. Calcium has many downstream signaling partners and targets, including many associated with actin dynamics. Due to the various membrane fluxes that are dependent on calcium, we do not include any direct calcium buffers or downstream targets other than cofilin. To couple with our mechanical model (see Section S2), we include cofilin and activated cofilin in the cytoplasm. Please note that we treat drebin and actin as constant values that serve to influence actin dynamics in the mechanical model, and therefore do not explicitly model their reaction-diffusion dynamics.

#### S2.7 Endoplasmic reticulum membrane (ER membrane)

We assume that there are IP_3_R along the whole ER membrane that bind IP_3_ and release Ca^2+^ into the cytoplasm. We also assume SERCA pumps reside on the whole ER membrane and pump Ca^2+^ back into the ER to replenish the ER calcium stores. Note that our flux boundary conditions for IP_3_R and SERCA pumps do not explicitly depend on the receptor and pump density.

#### S2.8 Endoplasmic reticulum (ER)

We model Ca^2+^ in the ER and include boundary flux reactions for flux through IP_3_Rs.

The diffusion rates for all species are in Table S2, all signaling reactions are listed in Table S3, and all reactions for each species is listed in Table S4.

#### S2.9 Geometric parameters used in the model

We model a single PC12 cell as a sphere of radius 5 m, with a nucleus of radius 2 m acting as an excluded volume in the middle of the cell soma. We include a single neurite of length 30 m and radius 1.25 m. Within this neurite, we model an endoplasmic reticulum as a cylinder of radius 0.25 m and length 25 m. The cytoplasm and plasma membrane have a volume-to-surface area ratio of n_PM_ = 1.12×10^-6^ m. The endoplasmic reticulum and ER membrane have a volume-to-surface area ratio of n_ER_ = ∼2×10^-7^ m. We use a 2D axisymmetric spatial model of a characteristic PC12 cell with a single neurite of interest. Geometric parameters are summarized in Table S5.

**Table S1.**
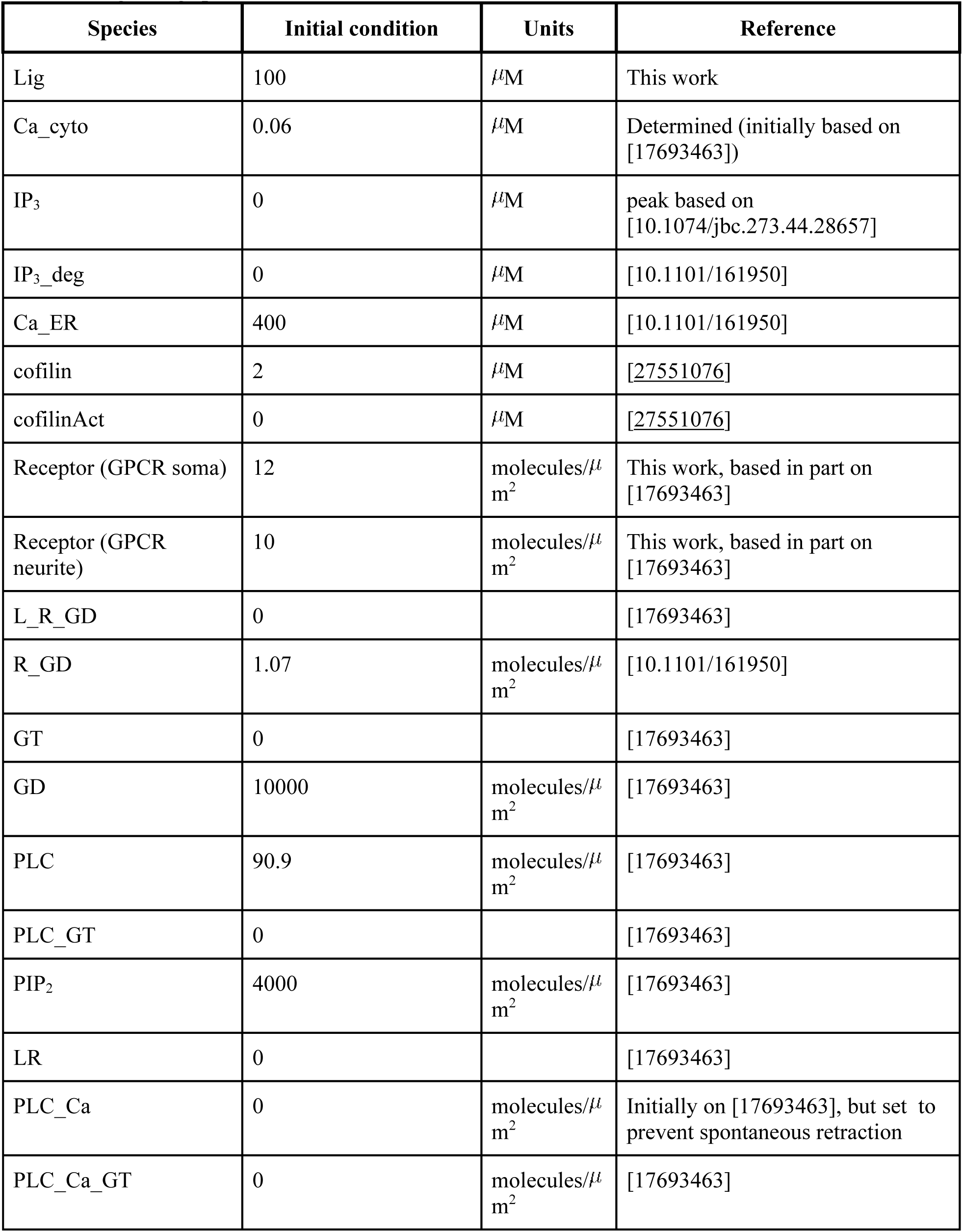
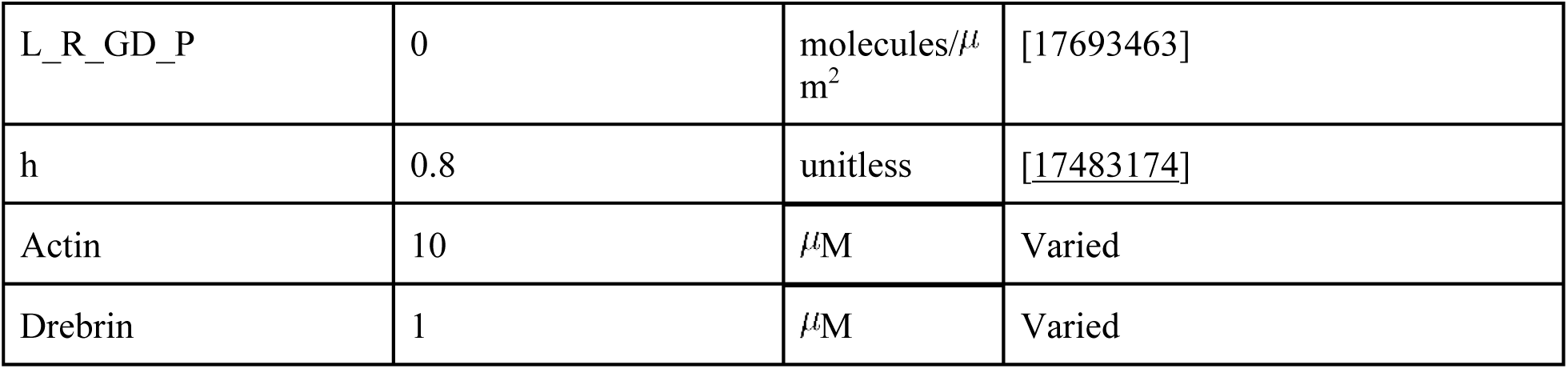
Signaling species and initial conditions.

**Table S2.**
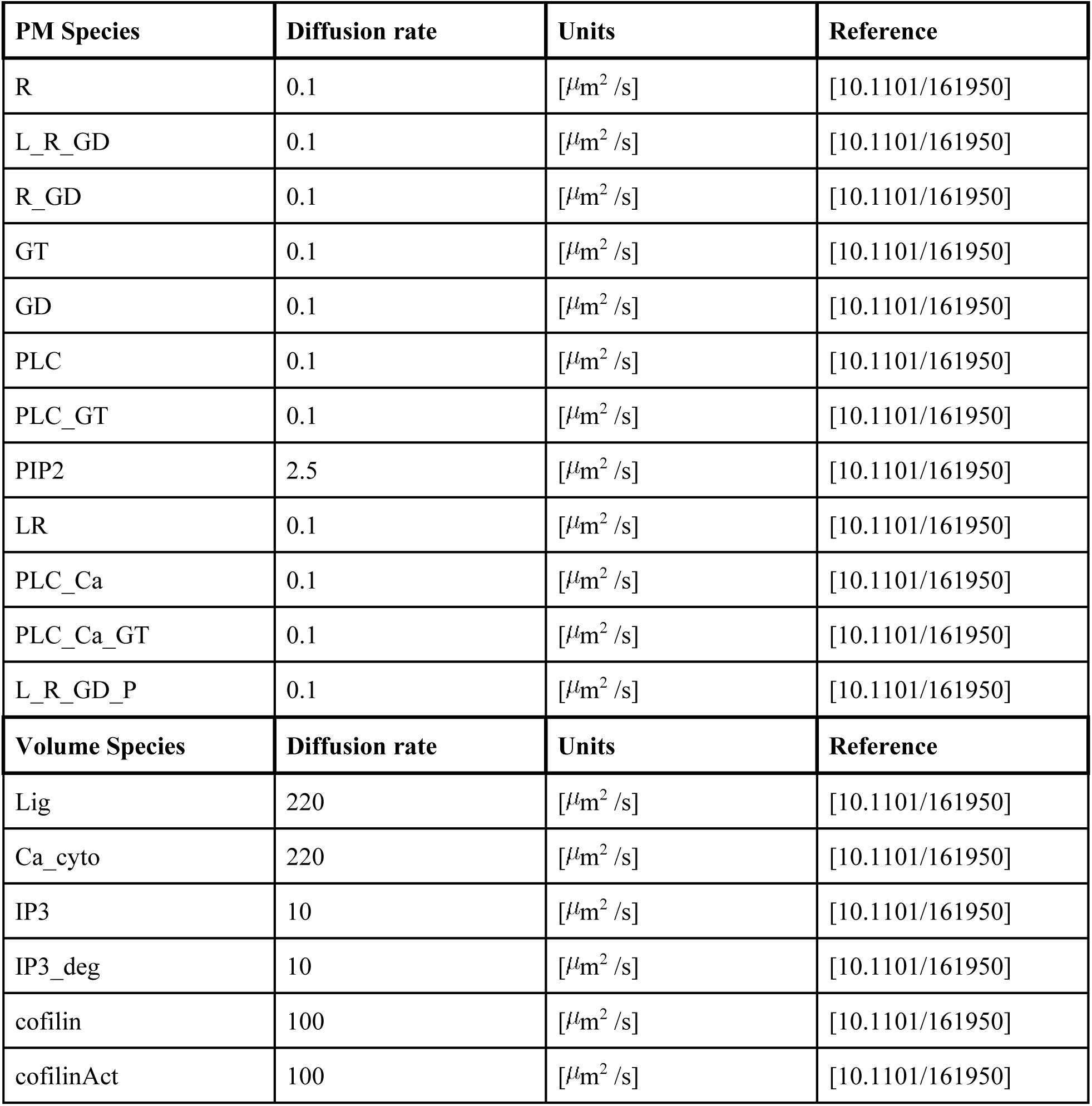
Diffusion rates for membrane and cytosolic proteins.

**Table S3.**
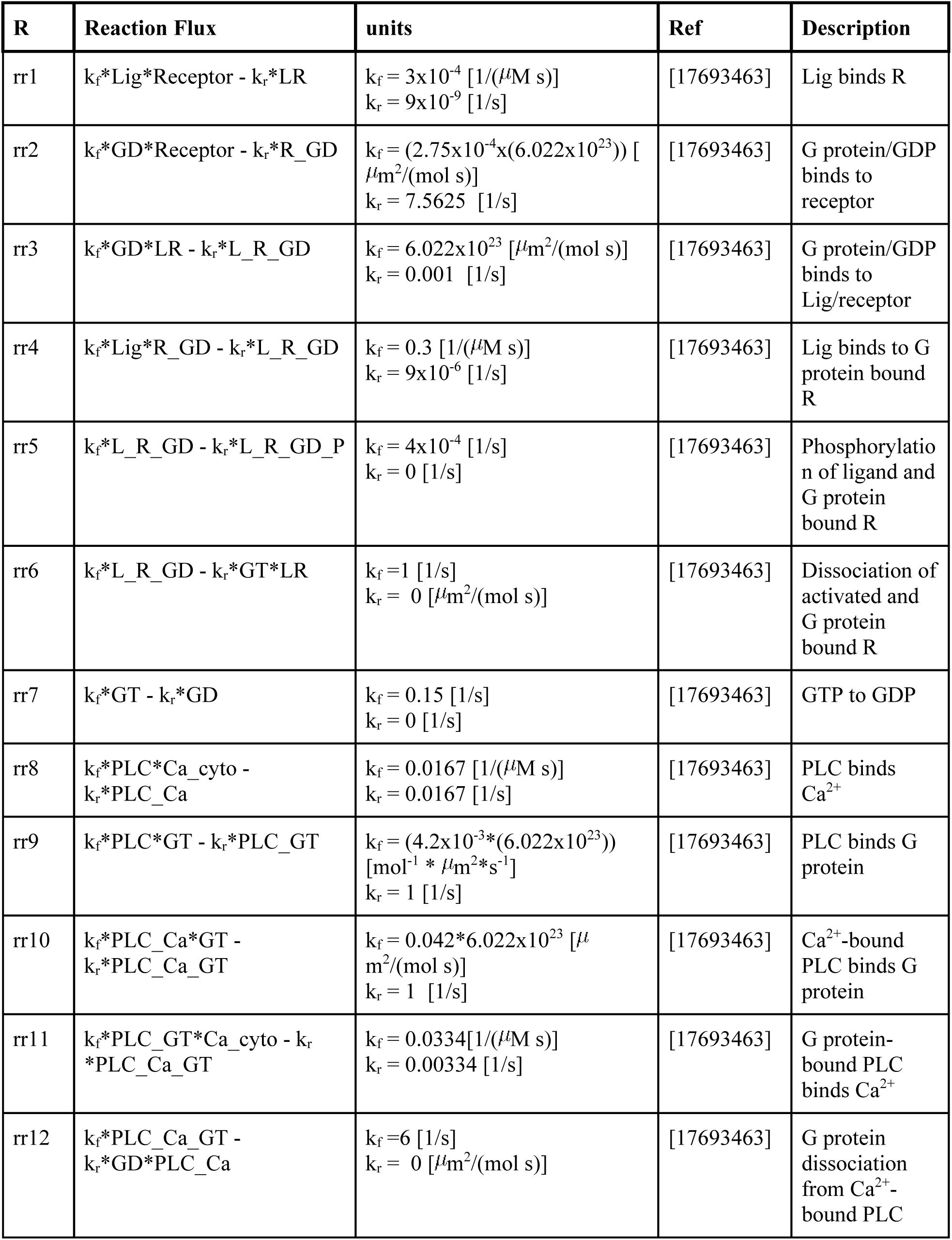
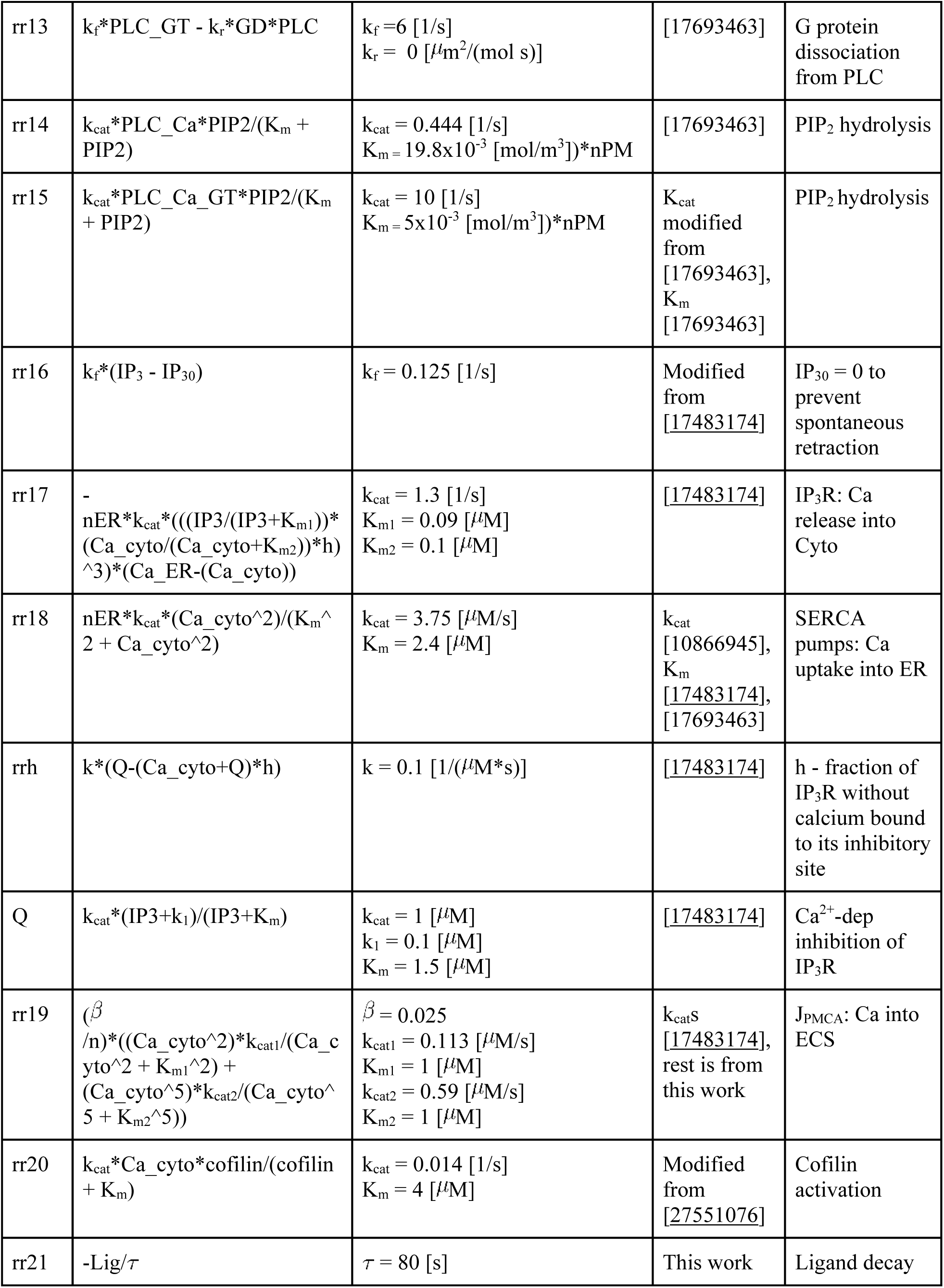
Reaction Rates.

| R | Reaction Flux | units | Ref | Description |
| --- | --- | --- | --- | --- |
| rr1 | $k_f \text{Lig} \cdot \text{Receptor} - k_r \text{LR}$ | $k_f = 3 \times 10^{-4} \text{ [1/(\mu M s)]}$<br>$k_r = 9 \times 10^{-9} \text{ [1/s]}$ | [17693463] | Lig binds R |
| rr2 | $k_f \text{GD} \cdot \text{Receptor} - k_r \text{R\_GD}$ | $k_f = (2.75 \times 10^{-4} \times (6.022 \times 10^{23})) \text{ [}\mu\text{m}^2/(\text{mol s})]$<br>$k_r = 7.5625 \text{ [1/s]}$ | [17693463] | G protein/GDP binds to receptor |
| rr3 | $k_f \text{GD} \cdot \text{LR} - k_r \text{L\_R\_GD}$ | $k_f = 6.022 \times 10^{23} \text{ [}\mu\text{m}^2/(\text{mol s})]$<br>$k_r = 0.001 \text{ [1/s]}$ | [17693463] | G protein/GDP binds to Lig/receptor |
| rr4 | $k_f \text{Lig} \cdot \text{R\_GD} - k_r \text{L\_R\_GD}$ | $k_f = 0.3 \text{ [1/(\mu M s)]}$<br>$k_r = 9 \times 10^{-6} \text{ [1/s]}$ | [17693463] | Lig binds to G protein bound R |
| rr5 | $k_f \text{L\_R\_GD} - k_r \text{L\_R\_GD\_P}$ | $k_f = 4 \times 10^{-4} \text{ [1/s]}$<br>$k_r = 0 \text{ [1/s]}$ | [17693463] | Phosphorylation of ligand and G protein bound R |
| rr6 | $k_f \text{L\_R\_GD} - k_r \text{GT} \cdot \text{LR}$ | $k_f = 1 \text{ [1/s]}$<br>$k_r = 0 \text{ [}\mu\text{m}^2/(\text{mol s})]$ | [17693463] | Dissociation of activated and G protein bound R |
| rr7 | $k_f \text{GT} - k_r \text{GD}$ | $k_f = 0.15 \text{ [1/s]}$<br>$k_r = 0 \text{ [1/s]}$ | [17693463] | GTP to GDP |
| rr8 | $k_f \text{PLC} \cdot \text{Ca}_{\text{cyto}} - k_r \text{PLC\_Ca}$ | $k_f = 0.0167 \text{ [1/(\mu M s)]}$<br>$k_r = 0.0167 \text{ [1/s]}$ | [17693463] | PLC binds $\text{Ca}^{2+}$ |
| rr9 | $k_f \text{PLC} \cdot \text{GT} - k_r \text{PLC\_GT}$ | $k_f = (4.2 \times 10^{-3} \times (6.022 \times 10^{23})) \text{ [mol}^{-1} \cdot \mu\text{m}^2 \cdot \text{s}^{-1}]$<br>$k_r = 1 \text{ [1/s]}$ | [17693463] | PLC binds G protein |
| rr10 | $k_f \text{PLC\_Ca} \cdot \text{GT} - k_r \text{PLC\_Ca\_GT}$ | $k_f = 0.042 \times 6.022 \times 10^{23} \text{ [}\mu\text{m}^2/(\text{mol s})]$<br>$k_r = 1 \text{ [1/s]}$ | [17693463] | $\text{Ca}^{2+}$-bound PLC binds G protein |
| rr11 | $k_f \text{PLC\_GT} \cdot \text{Ca}_{\text{cyto}} - k_r \text{PLC\_Ca\_GT}$ | $k_f = 0.0334 \text{ [1/(\mu M s)]}$<br>$k_r = 0.00334 \text{ [1/s]}$ | [17693463] | G protein-bound PLC binds $\text{Ca}^{2+}$ |
| rr12 | $k_f \text{PLC\_Ca\_GT} - k_r \text{GD} \cdot \text{PLC\_Ca}$ | $k_f = 6 \text{ [1/s]}$<br>$k_r = 0 \text{ [}\mu\text{m}^2/(\text{mol s})]$ | [17693463] | G protein dissociation from $\text{Ca}^{2+}$-bound PLC |
| rr13 | $k_f * \text{PLC\_GT} - k_r * \text{GD} * \text{PLC}$ | $k_f = 6 \text{ [1/s]}$<br>$k_r = 0 \text{ [}\mu\text{m}^2/(\text{mol s})]$ | [17693463] | G protein dissociation from PLC |
| rr14 | $k_{cat} * \text{PLC\_Ca} * \text{PIP}_2 / (K_m + \text{PIP}_2)$ | $k_{cat} = 0.444 \text{ [1/s]}$<br>$K_m = 19.8 \times 10^{-3} \text{ [mol/m}^3\text{)]} * n\text{PM}$ | [17693463] | PIP<sub>2</sub> hydrolysis |
| rr15 | $k_{cat} * \text{PLC\_Ca\_GT} * \text{PIP}_2 / (K_m + \text{PIP}_2)$ | $k_{cat} = 10 \text{ [1/s]}$<br>$K_m = 5 \times 10^{-3} \text{ [mol/m}^3\text{)]} * n\text{PM}$ | $K_{cat}$ modified from [17693463],<br>$K_m$ [17693463] | PIP<sub>2</sub> hydrolysis |
| rr16 | $k_f * (\text{IP}_3 - \text{IP}_{30})$ | $k_f = 0.125 \text{ [1/s]}$ | Modified from [17483174] | IP<sub>30</sub> = 0 to prevent spontaneous retraction |
| rr17 | $-n\text{ER} * k_{cat} * (((\text{IP}_3 / (\text{IP}_3 + K_{m1})) * (\text{Ca\_cyto} / (\text{Ca\_cyto} + K_{m2})) * h)^3 * (\text{Ca\_ER} - (\text{Ca\_cyto})))$ | $k_{cat} = 1.3 \text{ [1/s]}$<br>$K_{m1} = 0.09 \text{ [}\mu\text{M]}$<br>$K_{m2} = 0.1 \text{ [}\mu\text{M]}$ | [17483174] | IP<sub>3</sub>R: Ca release into Cyto |
| rr18 | $n\text{ER} * k_{cat} * (\text{Ca\_cyto}^2) / (K_m^2 + \text{Ca\_cyto}^2)$ | $k_{cat} = 3.75 \text{ [}\mu\text{M/s]}$<br>$K_m = 2.4 \text{ [}\mu\text{M]}$ | $k_{cat}$ [10866945],<br>$K_m$ [17483174], [17693463] | SERCA pumps: Ca uptake into ER |
| rrh | $k * (Q - (\text{Ca\_cyto} + Q) * h)$ | $k = 0.1 \text{ [1/(}\mu\text{M*s})]$ | [17483174] | h - fraction of IP<sub>3</sub>R without calcium bound to its inhibitory site |
| Q | $k_{cat} * (\text{IP}_3 + k_1) / (\text{IP}_3 + K_m)$ | $k_{cat} = 1 \text{ [}\mu\text{M]}$<br>$k_1 = 0.1 \text{ [}\mu\text{M]}$<br>$K_m = 1.5 \text{ [}\mu\text{M]}$ | [17483174] | Ca<sup>2+</sup>-dep inhibition of IP<sub>3</sub>R |
| rr19 | $(\beta / n) * ((\text{Ca\_cyto}^2) * k_{cat1} / (\text{Ca\_cyto}^2 + K_{m1}^2) + (\text{Ca\_cyto}^5) * k_{cat2} / (\text{Ca\_cyto}^5 + K_{m2}^5))$ | $\beta = 0.025$<br>$k_{cat1} = 0.113 \text{ [}\mu\text{M/s]}$<br>$K_{m1} = 1 \text{ [}\mu\text{M]}$<br>$k_{cat2} = 0.59 \text{ [}\mu\text{M/s]}$<br>$K_{m2} = 1 \text{ [}\mu\text{M]}$ | $k_{cat}$S [17483174], rest is from this work | J<sub>PMCA</sub>: Ca into ECS |
| rr20 | $k_{cat} * \text{Ca\_cyto} * \text{cofilin} / (\text{cofilin} + K_m)$ | $k_{cat} = 0.014 \text{ [1/s]}$<br>$K_m = 4 \text{ [}\mu\text{M]}$ | Modified from [27551076] | Cofilin activation |
| rr21 | $-\text{Lig} / \tau$ | $\tau = 80 \text{ [s]}$ | This work | Ligand decay |

**Table S4.**
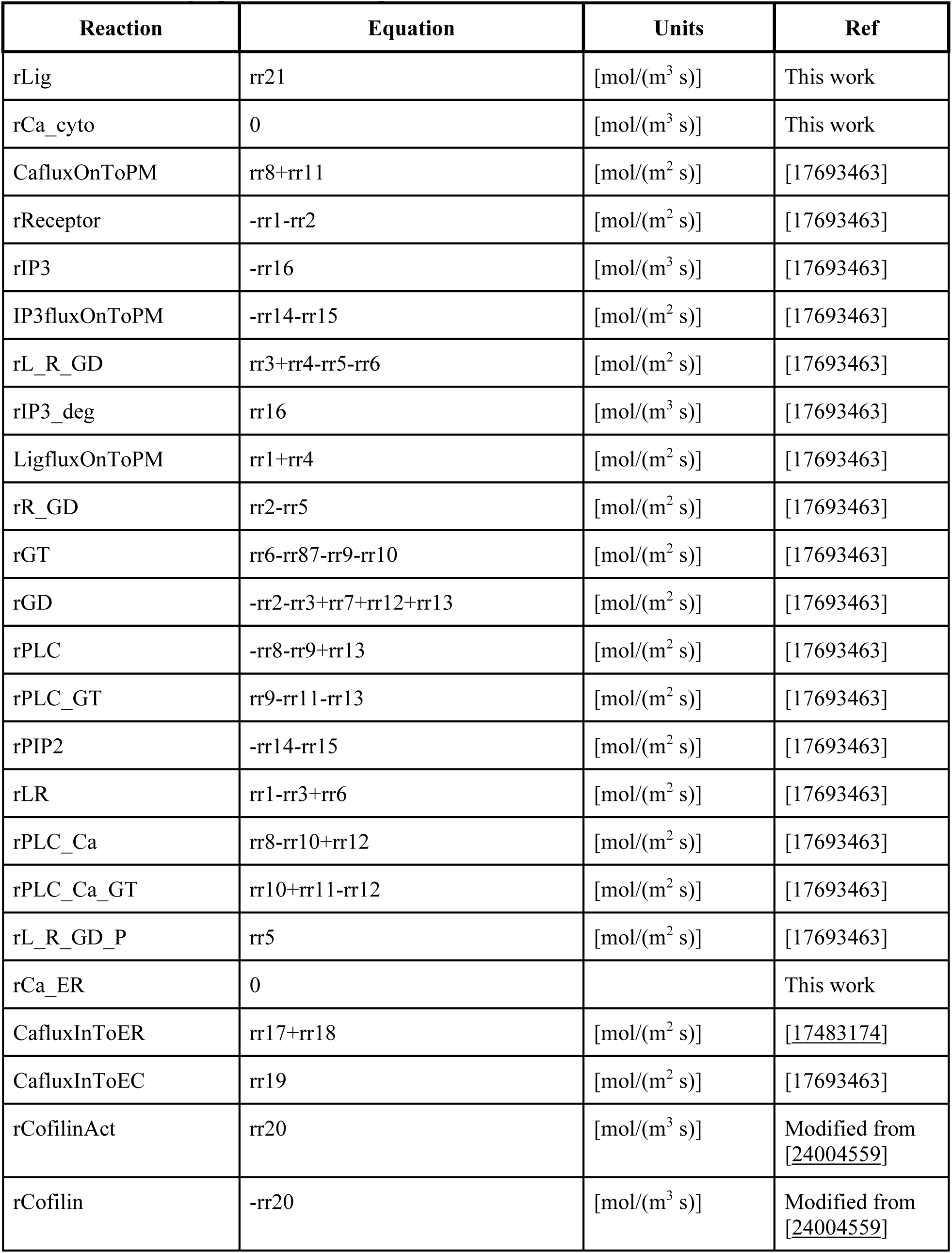
Governing equations for the species.

**Table S5.**
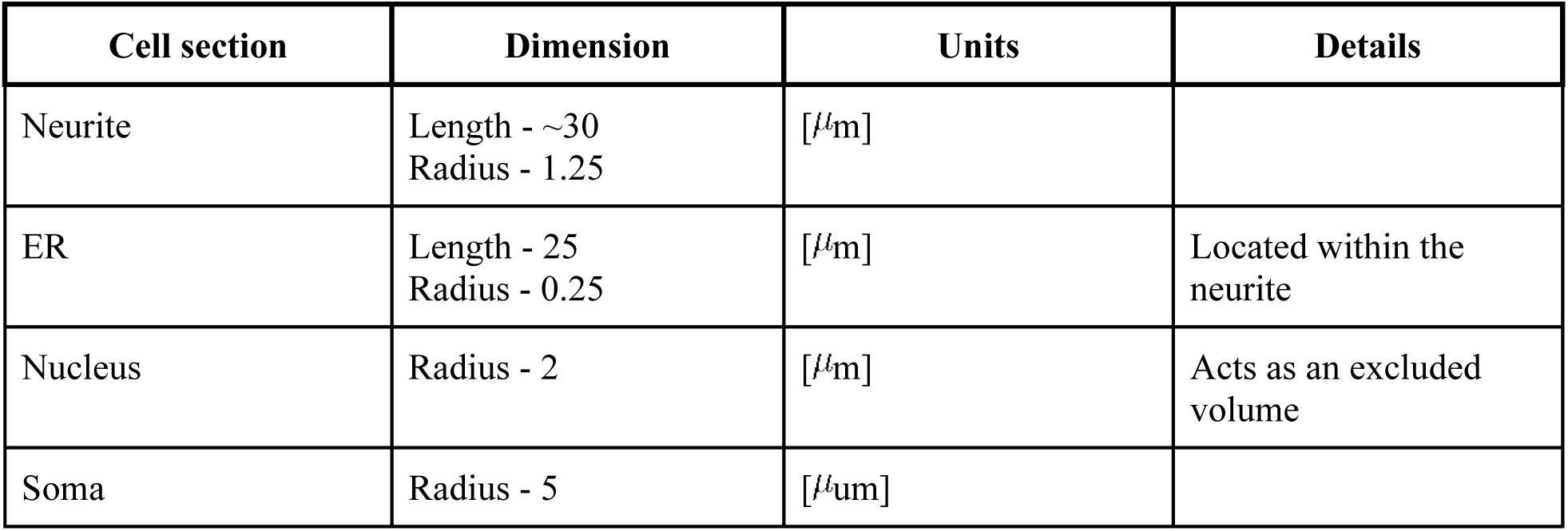
Geometric parameters in the spatial model.

### S3. Mechanical model

We developed an ordinary differential equation model of actin- and tension-driven mechanics controlling neurite retraction. Signaling dynamics are coupled and feed into the mechanical model [<u>19787029</u>] as spatially-averaged values. We based the mechanical model framework on experimental observations and existing literature [<u>24004559</u>, <u>20159147, 16679410</u>]. Using a framework proposed in [<u>24004559</u>] and experimental observations that neurite retraction as proportional to length, we developed a force balance written as:

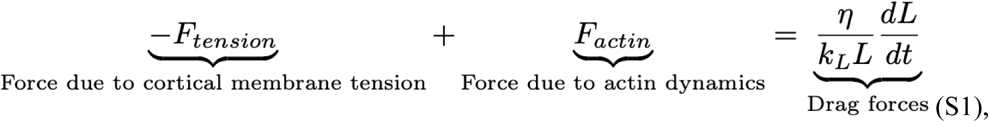

Where

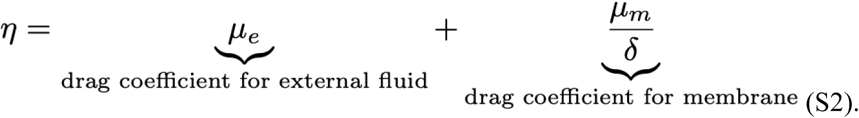

We arranging (Eq. S1), we find that the rate of neurite retraction could be given as

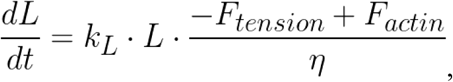

Where

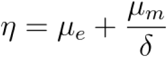

is the drag due to both the external fluid and membrane [<u>24004559</u>]. F_tension_ and F_actin_ have dependency on signaling input. In particular, F_tension_ is a function of tension, τ which itself depends on PIP_2_. F_actin_ is dependent on activated cofilin, and set amounts of total cofilin, drebrin, and actin. We can summarize the signaling inputs as τ = *f*(*PIP2*) and *F_actin_* = *g(cofilinACT)*

#### S3.1 Force due to tension

The tension force is directly proportional to the tension and is given by

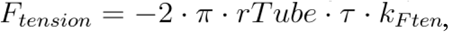

Where τ is tension [<u>24004559</u>]. Tension can be thought of as a Lagrange multiplier for conserved area, and tension change has three driving components regarding area change. We construct the model so that a decrease in area in the neurite, or an increased difference in area change between the neurite and base of the neurite causes an increase in tension. The rate of area change is given by

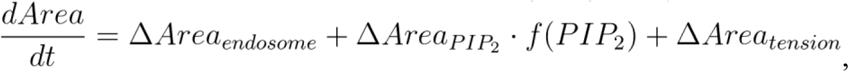

Where ΔArea represents the rate of change in area due to each contribution. Area change is due to the internalization of activated and phosphorylated GPCRs in endosomes, the decrease in surface area due to PIP_2_ hydrolysis that leaves the smaller DAG molecule in the membrane, and the increase in tension.

#### S3.2 Area change due to endocytosis of GPCRs

When GPCRs are both bound to ligand and phosphorylated, they are endocytosed in endosomes. This endocytosis process removes membrane and thus reduces the surface area. This rate of membrane area change is given by

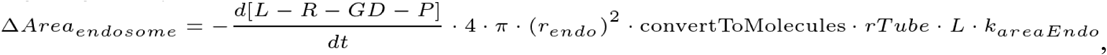

where r_endo_ is the radius of a typical endosome.

#### S3.3 Area change due to PIP_2_ hydrolysis

During PIP_2_ hydrolysis, PIP_2_ is converted into a cytosolic component, IP_3_, and a membrane bound component, DAG. DAG is smaller than PIP2, leading to a reduction in membrane surface area. This rate of membrane area change is given by

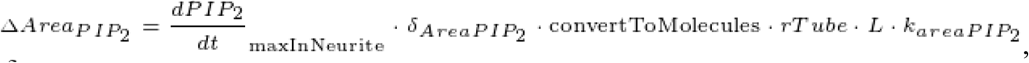

where δ_AreaPIP_2__ is the area difference between PIP_2_ size and DAG size. This area change is dependent on the rate of PIP_2_ hydrolysis. PIP_2_ dynamics have also been linked to effects on cytoskeleton, supporting the coupling of our model [12221130]. Note that in the total area change equation in S3.1, the PIP_2_ area term is multiplied by f(PIP_2_) which is a sigmoidal function dependent on the PIP_2_ density and is given by

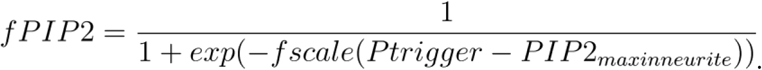

The sigmoidal form is based on [29473547].

#### S3.4 Area change due to tension

There is also a membrane area change due to tension. This term serves as feedback into the tension force equation because as tension changes due to area change, it can affect the rate of area change itself. This rate of membrane area change is based on [<u>28648660</u>] and is given by

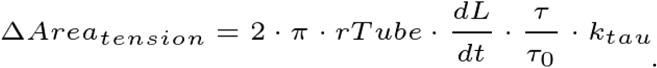

τ_0_ is the initial tension value.

Combining these effects, we get the follow equation for tension,

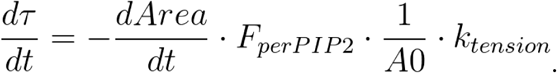

#### S3.5 Actin force

The actin force is influenced by various actin-related proteins. We note that actin dynamics are highly complicated quite extensively studied [24209839, 10.1523/JNEUROSCI.11-07-01918.1991, 10.1016/0896-6273(88)90124-9, 20159147]. We also highlight that we experimentally observe unusual actin dynamics during this neurite retraction. Therefore, we present a phenomenological model for actin dynamics based on a stress-strain framework. We construct that model such that cofilin reduces the actin pushing force, and drebrin and actin slow the reduction in actin pushing force. Eventually the decrease in pushing force becomes a negative force that can be attributed to retrograde flow [10.1016/S0955-0674(97)80152-4, 24198333]. Therefore, actin force is given by

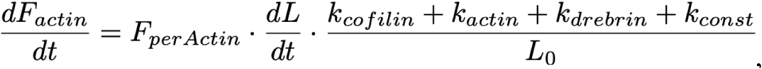

where the k contributions are from cofilin [10.1002/(SICI)1097-0169(1998)39:2<172::AID-CM8>3.0.CO;2-8], actin [10.1523/JNEUROSCI.11-07-01918.1991], and drebrin [<u>21175132,23696644</u>].

The k equations are given by

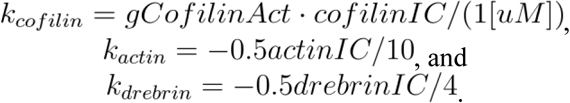

The cofilin contribution is again dependent on a sigmoidal function [29473547] that is dependent on activated cofilin (cofilinAct), and is given by

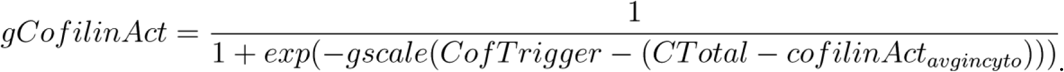

The various parameters in the mechanical model are given in Table S6.

**Table S6.**
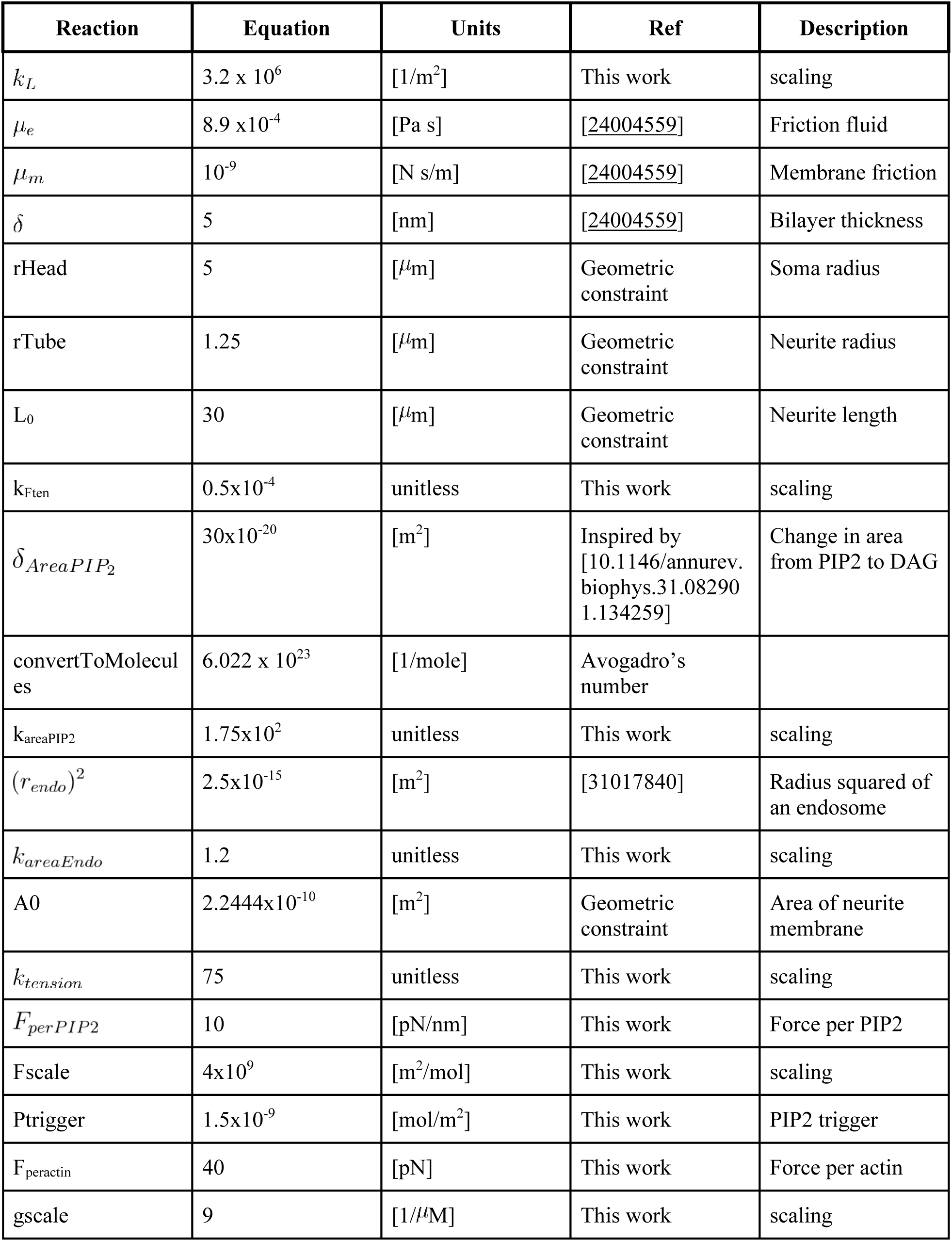
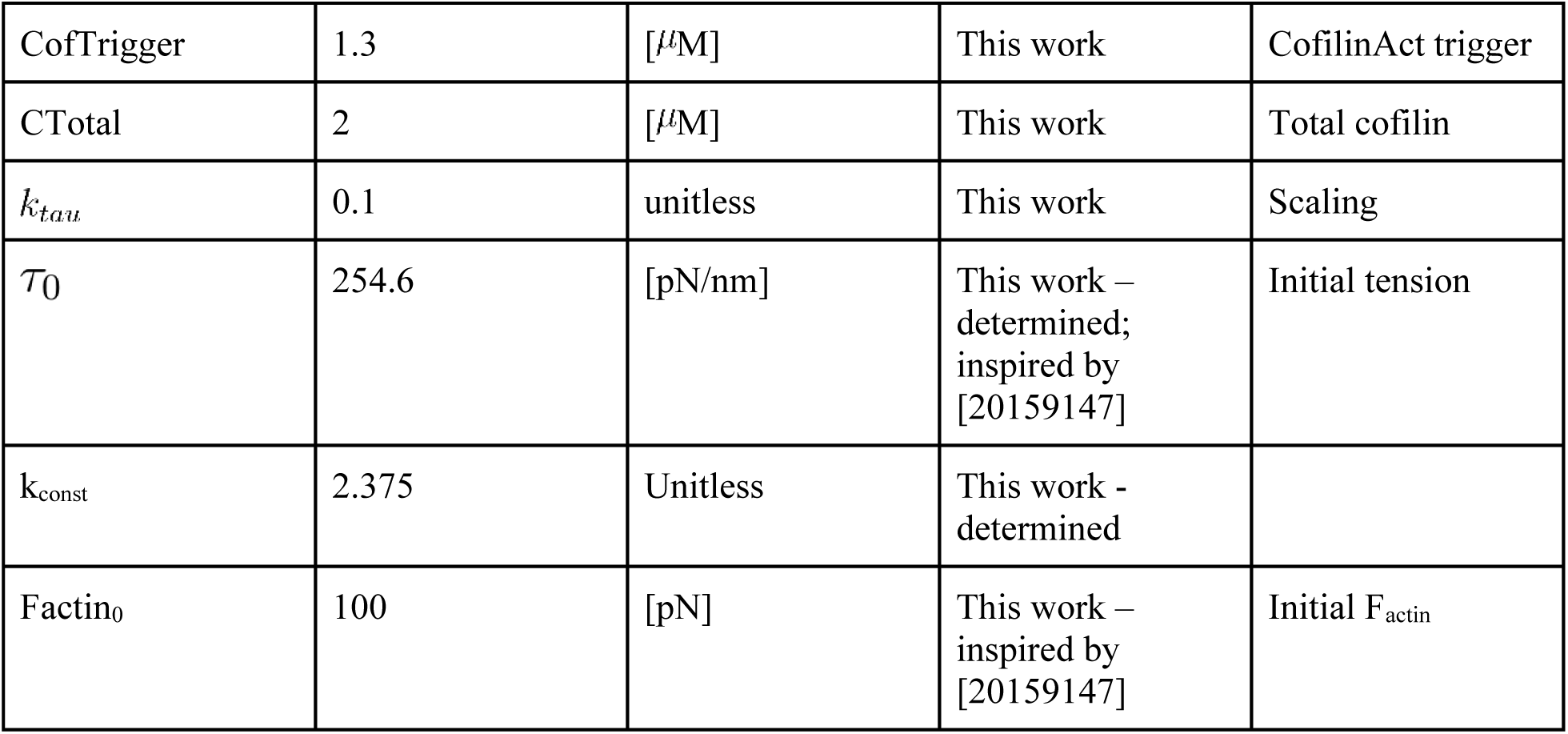
Parameters in the mechanical model.

#### S3.6 Overexpression trials

To capture protein overexpression experiments for actin, cofilin, and drebrin, the concentration of overexpressed protein was input into the corresponding protein’s k equation. For example, actin is believed to be overexpressed from about 10 M to 30 M, so actinIC goes from 10 to 30 in actin’s k equation. In this way, we capture the effects of protein overexpression on actin force dynamics. See Table S7 for the estimated protein concentrations during each overexpression trial.

**Table S7.**
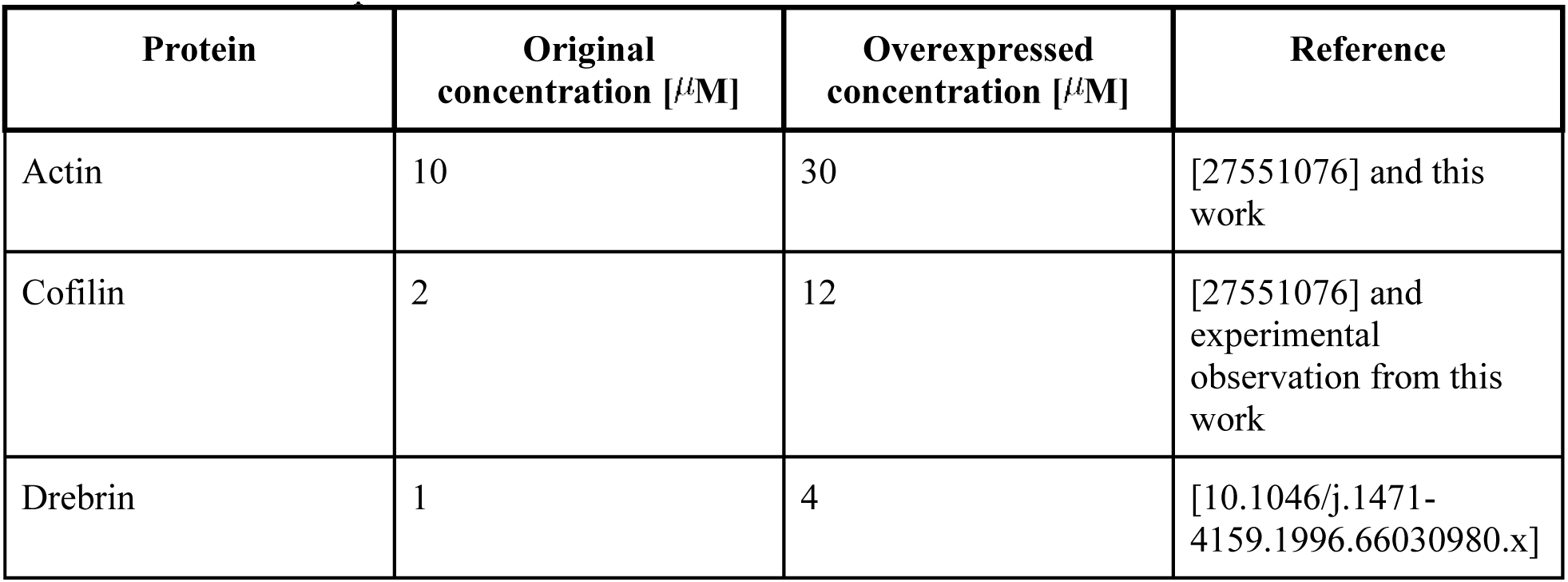
Actin-related protein concentrations.

#### S3.7 Hyperosmotic stress model

To elicit the role of tension in neurite retraction dynamics, hyperosmotic stress experiments were conducted. During these trials, neurite retraction was observed in the absence of ligand stimulation, confirming the role of tension in retraction. To model hyperosmotic conditions, we assumed that that hyperosmotic environment will trigger volume reduction and an initial increase in membrane tension. Therefore, we increase our initial tension value in the absence of ligand input.

## s4. Western Blots to assess protein levels used in modeling

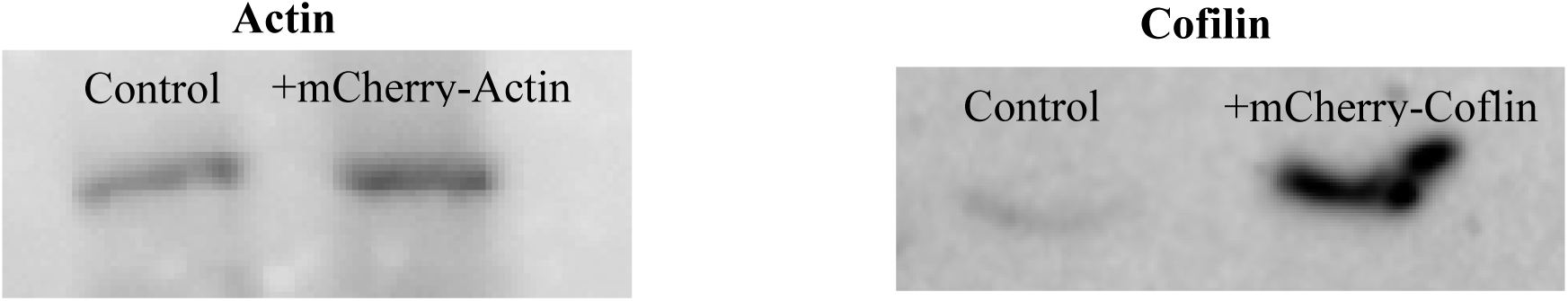

V1-2 (attached) Videos showing neurite retraction with carbachol stimulation seen for PC12 cells labeled with Calcium Green (left) and DIC (right). The corresponding screen shots are shown in Fig 1 and below. Experimental details can be found in the text.

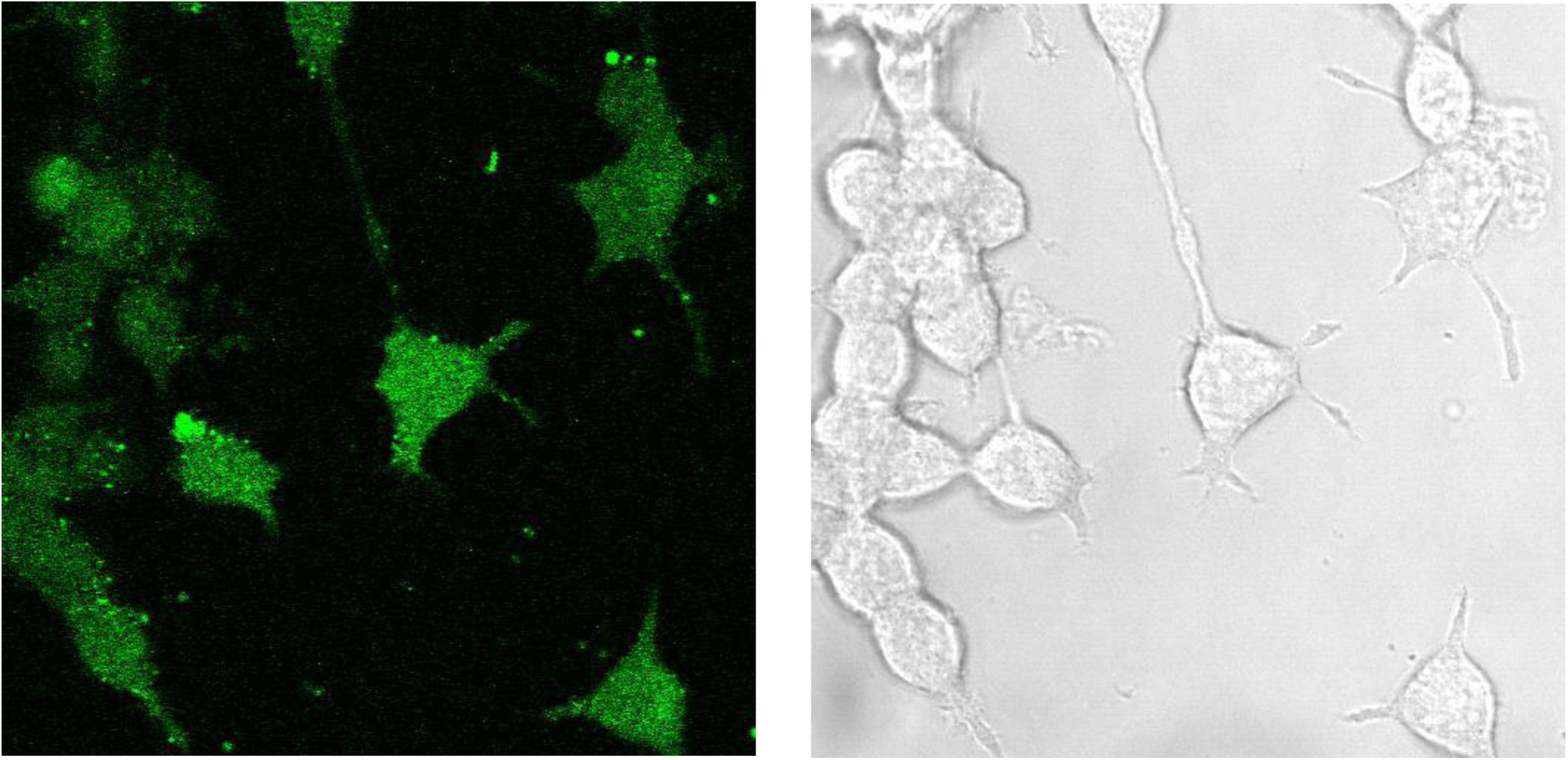

V3 (attached) Neurite retraction with carbachol stimulation as monitored by eCFP-B2R where the associated screen shots are shown below and can be found in Fig2 B-**C.** Experimental details can be found in the text.

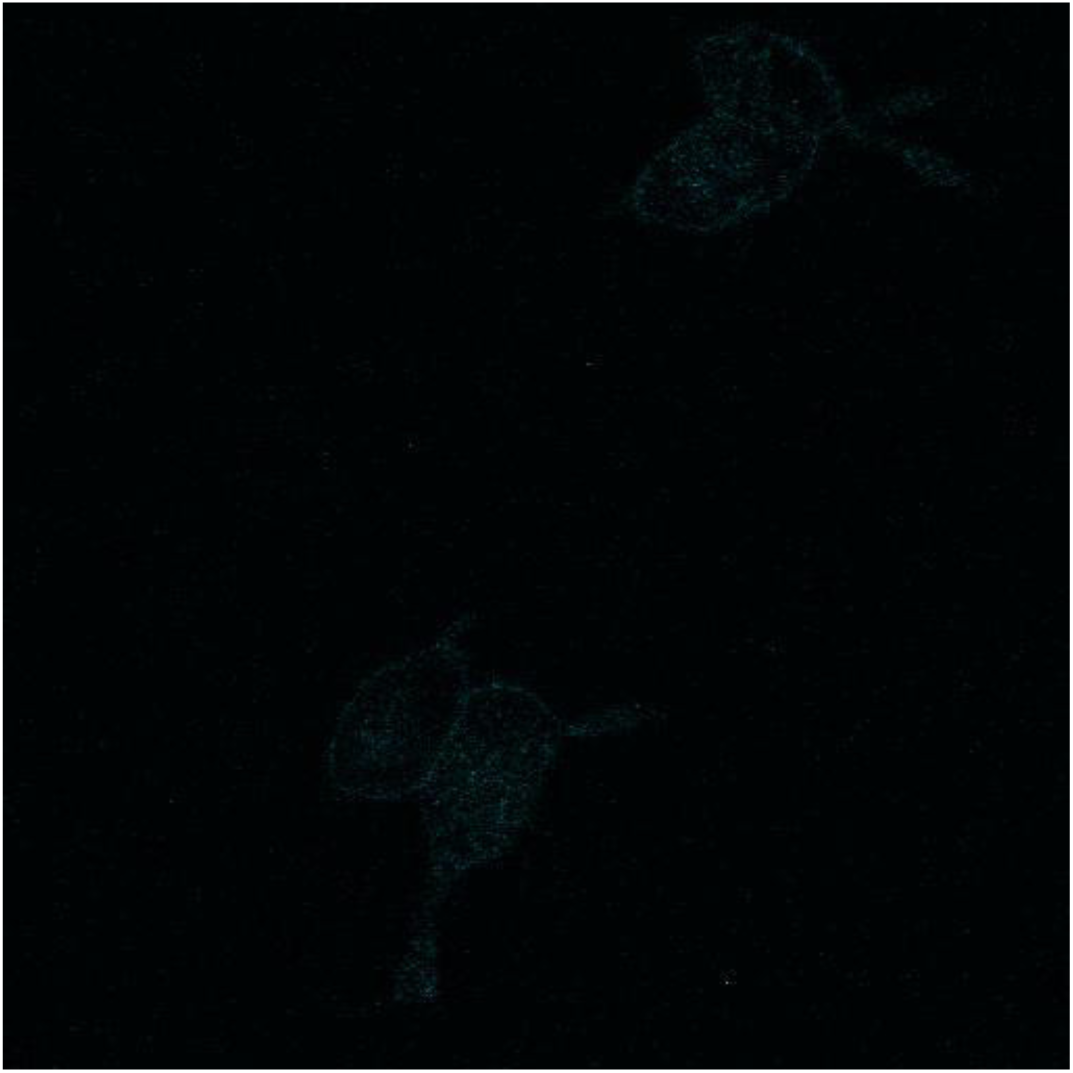

V4 (attached) The behavior of PI(4,5)P_2_ with carbachol-induced neurite retraction as followed by PH-PLCδ1 where the screen shots are shown below and in Fig2 F-G. Experimental details can be found in the text.

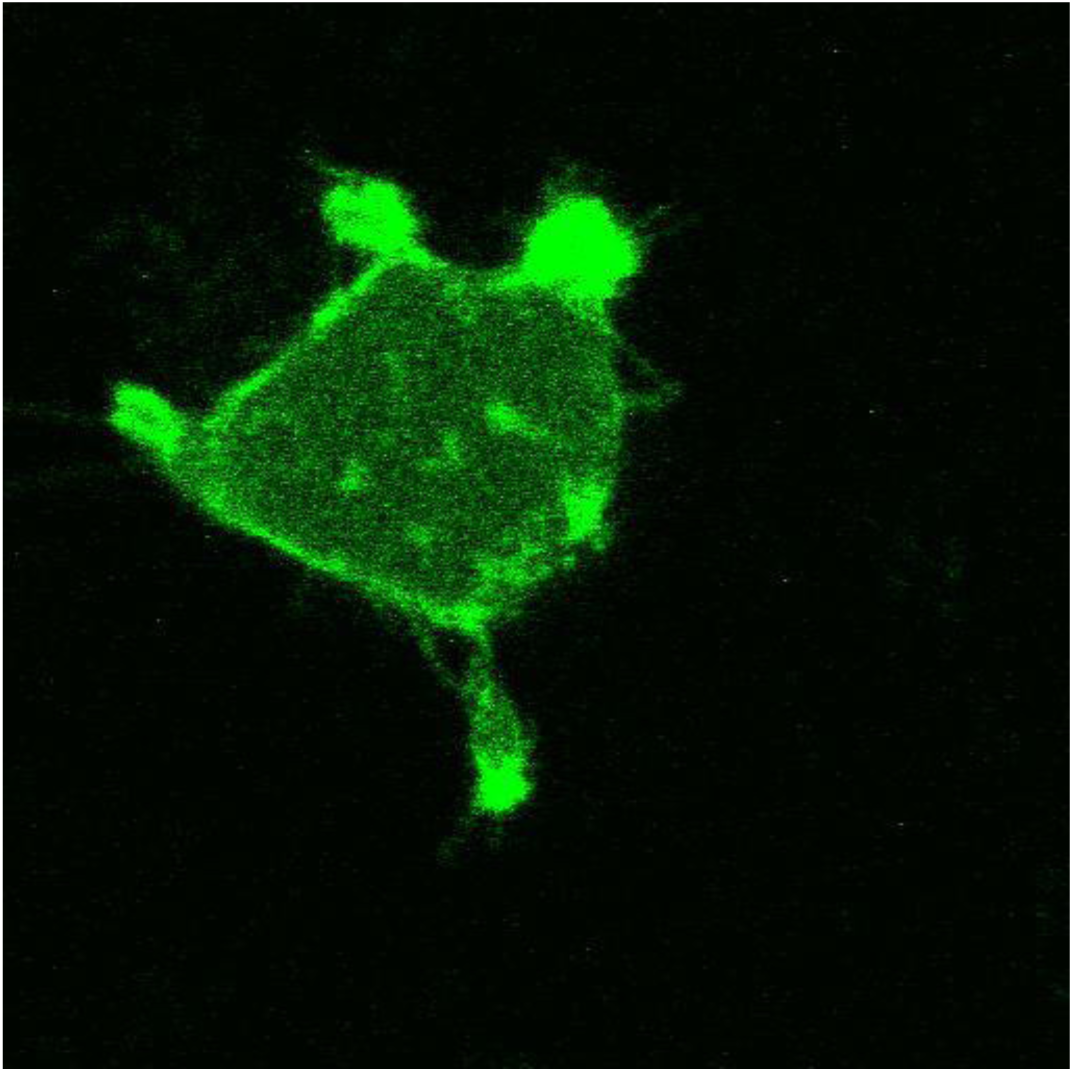

V5 (attached) The behavior of mCherry-actin with carbachol-induced neurite retraction where the screen shots are shown below and in Fig. 4 D, G, J. Experimental details can be found in the text.

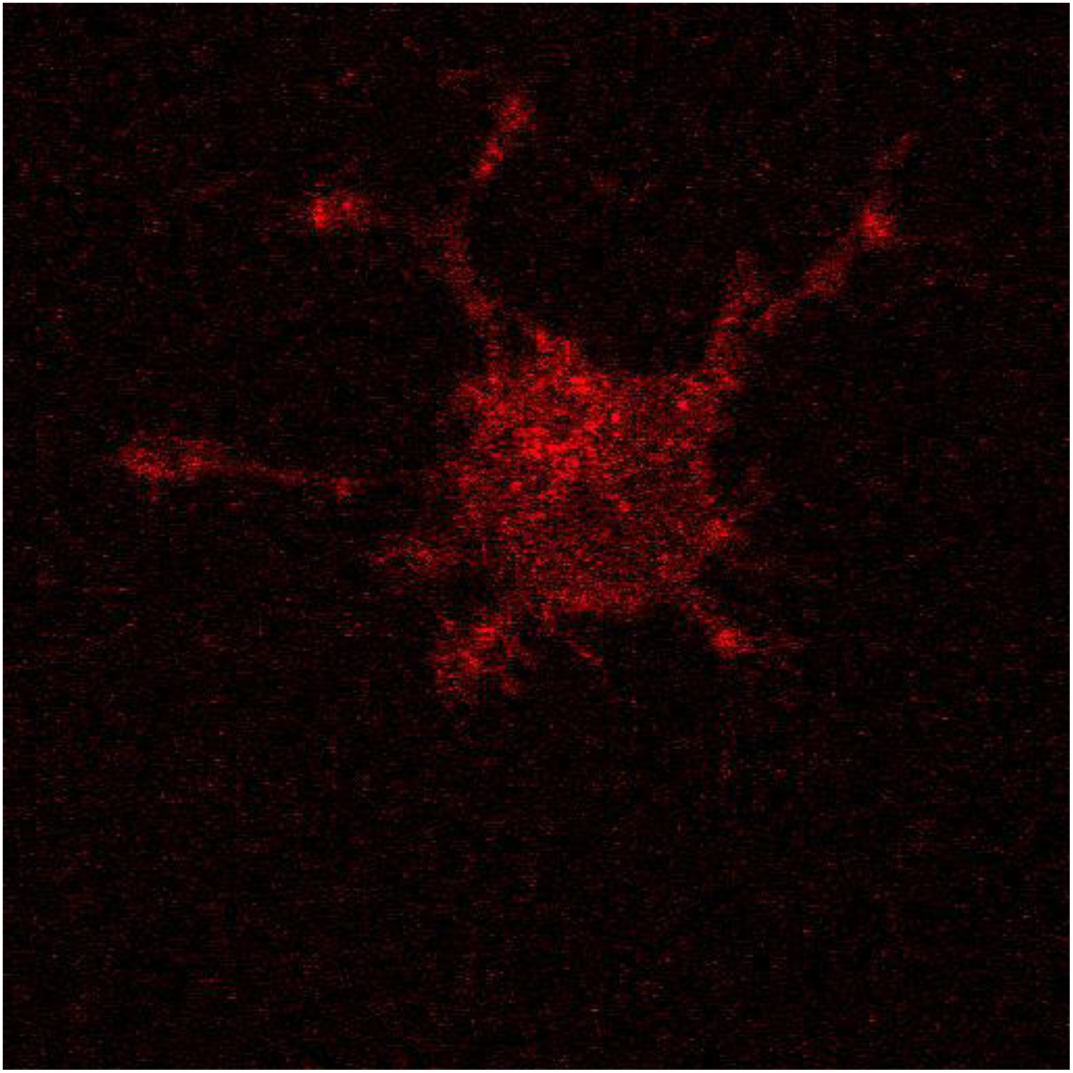

V6 (attached) The behavior of mCherry-cofilin with carbachol-induced neurite where the screen shots are shown below and in Fig4 E, H, K. Experimental details can be found in the text.

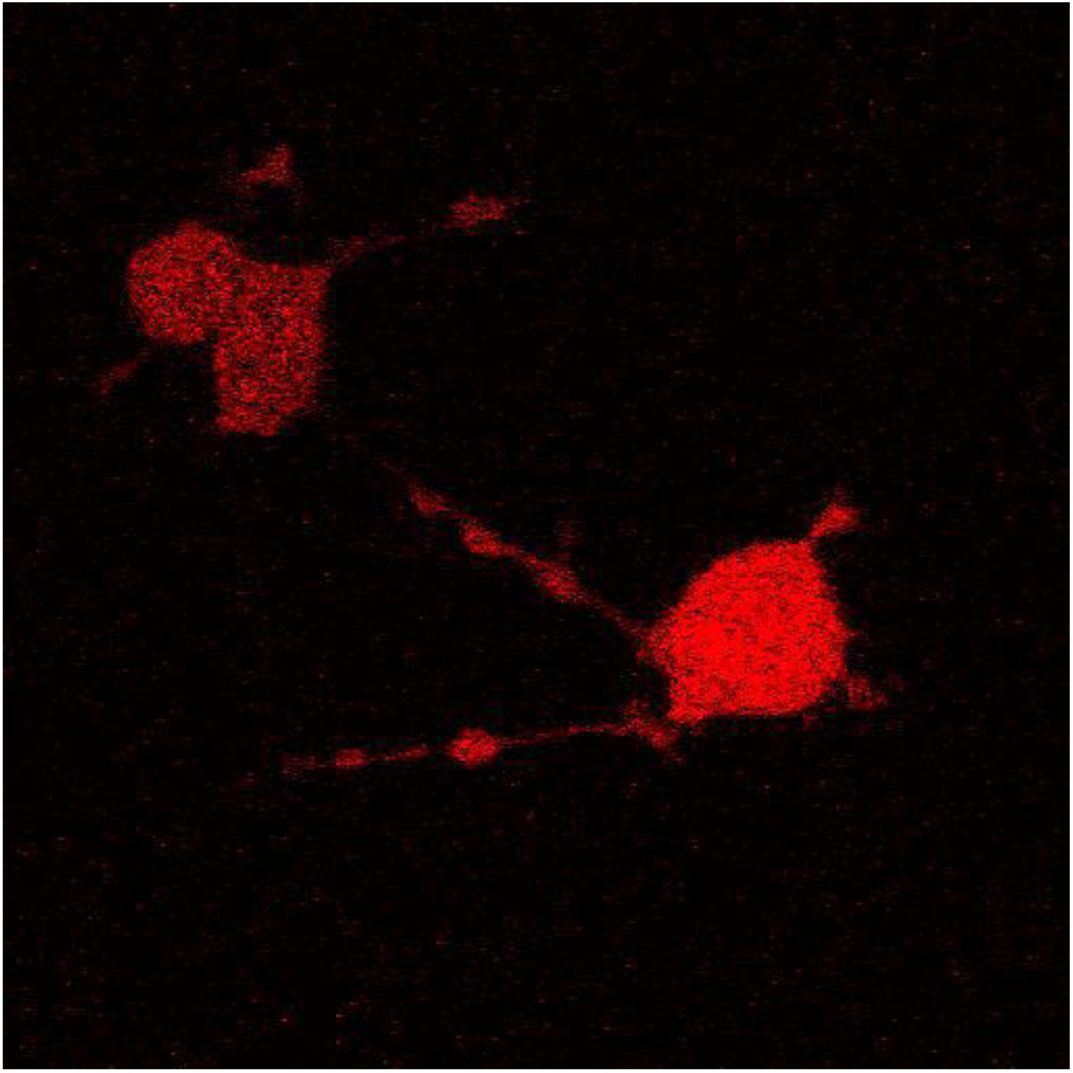

