## Supplementary material for "Gq-mediated calcium dynamics and membrane tension modulate neurite plasticity": Data associated with modeling and experimental data

$$\frac{\partial S_x}{\partial t} = D_{S_x} \nabla^2 S_x + f(S_x, ...),$$

where f($S_x$,...) are any reactions regarding species $S_x$. For volumetric species, flux into or out of the volume is given by a membrane flux boundary condition written as

**S2.9 Geometric parameters used in the model**
We model a single PC12 cell as a sphere of radius 5 $\mu$m, with a nucleus of radius 2$\mu$m acting as an excluded volume in the middle of the cell soma. We include a single neurite of length 30 $\mu$m and radius 1.25 $\mu$m. Within this neurite, we model an endoplasmic reticulum as a cylinder of radius 0.25 $\mu$m and length 25 $\mu$m. The cytoplasm and plasma membrane have a volume-to-surface area ratio of $n_{PM}$ = $1.12x10^{-6}$ m. The endoplasmic reticulum and ER membrane have a volume-to-surface area ratio of $n_{ER}$ = ~$2x10^{-7}$ m. We use a 2D axisymmetric spatial model of a characteristic PC12 cell with a single neurite of interest. Geometric parameters are summarized in Table S5.

**Table S1. Signaling species and initial conditions**

| Species | Initial condition | Units | Reference |
|---|---|---|---|
| Lig | 100 | $\mu$M | This work |
| Ca_cyto | 0.06 | $\mu$M | Determined (initially based on [17693463]) |
| $IP_3$ | 0 | $\mu$M | peak based on [10.1074/jbc.273.44.28657] |
| $IP_3$_deg | 0 | $\mu$M | [10.1101/161950] |
| Ca_ER | 400 | $\mu$M | [10.1101/161950] |
| cofilin | 2 | $\mu$M | [27551076] |
| cofilinAct | 0 | $\mu$M | [27551076] |
| Receptor (GPCR soma) | 12 | molecules/$\mu m^2$ | This work, based in part on [17693463] |
| Receptor (GPCR neurite) | 10 | molecules/$\mu m^2$ | This work, based in part on [17693463] |
| L_R_GD | 0 | | [17693463] |
| R_GD | 1.07 | molecules/$\mu m^2$ | [10.1101/161950] |
| GT | 0 | | [17693463] |
| GD | 10000 | molecules/$\mu m^2$ | [17693463] |
| PLC | 90.9 | molecules/$\mu m^2$ | [17693463] |
| PLC_GT | 0 | | [17693463] |
| $PIP_2$ | 4000 | molecules/$\mu m^2$ | [17693463] |
| LR | 0 | | [17693463] |
| PLC_Ca | 0 | molecules/$\mu m^2$ | Initially on [17693463], but set to prevent spontaneous retraction |
| PLC_Ca_GT | 0 | molecules/$\mu m^2$ | [17693463] |

| | | | |
|---|---|---|---|
| L_R_GD_P | 0 | molecules/$\mu m^2$ | [17693463] |
| h | 0.8 | unitless | [17483174] |
| Actin | 10 | $\mu$M | Varied |
| Drebrin | 1 | $\mu$M | Varied |

**Table S2. Diffusion rates for membrane and cytosolic proteins**

| **PM Species** | **Diffusion rate** | **Units** | **Reference** |
|---|---|---|---|
| R | 0.1 | [$\mu m^2$ /s] | [10.1101/161950] |
| L_R_GD | 0.1 | [$\mu m^2$ /s] | [10.1101/161950] |
| R_GD | 0.1 | [$\mu m^2$ /s] | [10.1101/161950] |
| GT | 0.1 | [$\mu m^2$ /s] | [10.1101/161950] |
| GD | 0.1 | [$\mu m^2$ /s] | [10.1101/161950] |
| PLC | 0.1 | [$\mu m^2$ /s] | [10.1101/161950] |
| PLC_GT | 0.1 | [$\mu m^2$ /s] | [10.1101/161950] |
| PIP2 | 2.5 | [$\mu m^2$ /s] | [10.1101/161950] |
| LR | 0.1 | [$\mu m^2$ /s] | [10.1101/161950] |
| PLC_Ca | 0.1 | [$\mu m^2$ /s] | [10.1101/161950] |
| PLC_Ca_GT | 0.1 | [$\mu m^2$ /s] | [10.1101/161950] |
| L_R_GD_P | 0.1 | [$\mu m^2$ /s] | [10.1101/161950] |
| **Volume Species** | **Diffusion rate** | **Units** | **Reference** |
| Lig | 220 | [$\mu m^2$ /s] | [10.1101/161950] |
| Ca_cyto | 220 | [$\mu m^2$ /s] | [10.1101/161950] |
| IP3 | 10 | [$\mu m^2$ /s] | [10.1101/161950] |
| IP3_deg | 10 | [$\mu m^2$ /s] | [10.1101/161950] |
| cofilin | 100 | [$\mu m^2$ /s] | [10.1101/161950] |
| cofilinAct | 100 | [$\mu m^2$ /s] | [10.1101/161950] |

**Table S3. Reaction Rates**

| R | Reaction Flux | units | Ref | Description |
|---|---|---|---|---|
| rr1 | $k_f$*Lig*Receptor - $k_r$*LR | $k_f = 3x10^{-4}$ [1/($\mu$M s)]<br>$k_r = 9x10^{-9}$ [1/s] | [17693463] | Lig binds R |
| rr2 | $k_f$*GD*Receptor - $k_r$*R_GD | $k_f = (2.75x10^{-4}x(6.022x10^{23}))$ [$\mu m^2$/(mol s)]<br>$k_r$ = 7.5625 [1/s] | [17693463] | G protein/GDP binds to receptor |
| rr3 | $k_f$*GD*LR - $k_r$*L_R_GD | $k_f = 6.022x10^{23}$ [$\mu m^2$/(mol s)]<br>$k_r$ = 0.001 [1/s] | [17693463] | G protein/GDP binds to Lig/receptor |
| rr4 | $k_f$*Lig*R_GD - $k_r$*L_R_GD | $k_f$ = 0.3 [1/($\mu$M s)]<br>$k_r = 9x10^{-6}$ [1/s] | [17693463] | Lig binds to G protein bound R |
| rr5 | $k_f$*L_R_GD - $k_r$*L_R_GD_P | $k_f = 4x10^{-4}$ [1/s]<br>$k_r$ = 0 [1/s] | [17693463] | Phosphorylation of ligand and G protein bound R |
| rr6 | $k_f$*L_R_GD - $k_r$*GT*LR | $k_f$ =1 [1/s]<br>$k_r$ = 0 [$\mu m^2$/(mol s)] | [17693463] | Dissociation of activated and G protein bound R |
| rr7 | $k_f$*GT - $k_r$*GD | $k_f$ = 0.15 [1/s]<br>$k_r$ = 0 [1/s] | [17693463] | GTP to GDP |
| rr8 | $k_f$*PLC*Ca_cyto - $k_r$*PLC_Ca | $k_f$ = 0.0167 [1/($\mu$M s)]<br>$k_r$ = 0.0167 [1/s] | [17693463] | PLC binds $Ca^{2+}$ |
| rr9 | $k_f$*PLC*GT - $k_r$*PLC_GT | $k_f = (4.2x10^{-3}*(6.022x10^{23}))$ [$mol^{-1} * \mu m^2 * s^{-1}$]<br>$k_r$ = 1 [1/s] | [17693463] | PLC binds G protein |
| rr10 | $k_f$*PLC_Ca*GT - $k_r$*PLC_Ca_GT | $k_f = 0.042*6.022x10^{23}$ [$\mu m^2$/(mol s)]<br>$k_r$ = 1 [1/s] | [17693463] | $Ca^{2+}$-bound PLC binds G protein |
| rr11 | $k_f$*PLC_GT*Ca_cyto - $k_r$ *PLC_Ca_GT | $k_f$ = 0.0334[1/($\mu$M s)]<br>$k_r$ = 0.00334 [1/s] | [17693463] | G protein-bound PLC binds $Ca^{2+}$ |
| rr12 | $k_f$*PLC_Ca_GT - $k_r$*GD*PLC_Ca | $k_f$ =6 [1/s]<br>$k_r$ = 0 [$\mu m^2$/(mol s)] | [17693463] | G protein dissociation from $Ca^{2+}$-bound PLC |

| | | | | |
|---|---|---|---|---|
| rr13 | $k_f$*PLC_GT - $k_r$*GD*PLC | $k_f$ =6 [1/s]<br>$k_r$ = 0 [$\mu m^2$/(mol s)] | [17693463] | G protein dissociation from PLC |
| rr14 | $k_{cat}$*PLC_Ca*PIP2/($K_m$ + PIP2) | $k_{cat}$ = 0.444 [1/s]<br>$K_{m\,=}$ 19.8x$10^{-3}$ [mol/$m^3$])*nPM | [17693463] | $PIP_2$ hydrolysis |
| rr15 | $k_{cat}$*PLC_Ca_GT*PIP2/($K_m$ + PIP2) | $k_{cat}$ = 10 [1/s]<br>$K_{m\,=}$ 5x$10^{-3}$ [mol/$m^3$])*nPM | $K_{cat}$ modified from [17693463], $K_m$ [17693463] | $PIP_2$ hydrolysis |
| rr16 | $k_f$*($IP_3$ - $IP_{30}$) | $k_f$ = 0.125 [1/s] | Modified from [17483174] | $IP_{30}$ = 0 to prevent spontaneous retraction |
| rr17 | -nER*$k_{cat}$*(((IP3/(IP3+$K_{m1}$))*(Ca_cyto/(Ca_cyto+$K_{m2}$))*h)^3)*(Ca_ER-(Ca_cyto)) | $k_{cat}$ = 1.3 [1/s]<br>$K_{m1}$ = 0.09 [$\mu$M]<br>$K_{m2}$ = 0.1 [$\mu$M] | [17483174] | $IP_3$R: Ca release into Cyto |
| rr18 | nER*$k_{cat}$*(Ca_cyto^2)/($K_m$^2 + Ca_cyto^2) | $k_{cat}$ = 3.75 [$\mu$M/s]<br>$K_m$ = 2.4 [$\mu$M] | $k_{cat}$ [10866945], $K_m$ [17483174], [17693463] | SERCA pumps: Ca uptake into ER |
| rrh | k*(Q-(Ca_cyto+Q)*h) | k = 0.1 [1/($\mu$M*s)] | [17483174] | h - fraction of $IP_3$R without calcium bound to its inhibitory site |
| Q | $k_{cat}$*(IP3+$k_1$)/(IP3+$K_m$) | $k_{cat}$ = 1 [$\mu$M]<br>$k_1$ = 0.1 [$\mu$M]<br>$K_m$ = 1.5 [$\mu$M] | [17483174] | $Ca^{2+}$-dep inhibition of $IP_3$R |
| rr19 | ($\beta$/n)*((Ca_cyto^2)*$k_{cat1}$/(Ca_cyto^2 + $K_{m1}$^2) + (Ca_cyto^5)*$k_{cat2}$/(Ca_cyto^5 + $K_{m2}$^5)) | $\beta$ = 0.025<br>$k_{cat1}$ = 0.113 [$\mu$M/s]<br>$K_{m1}$ = 1 [$\mu$M]<br>$k_{cat2}$ = 0.59 [$\mu$M/s]<br>$K_{m2}$ = 1 [$\mu$M] | $k_{cat}$s [17483174], rest is from this work | $J_{PMCA}$: Ca into ECS |
| rr20 | $k_{cat}$*Ca_cyto*cofilin/(cofilin + $K_m$) | $k_{cat}$ = 0.014 [1/s]<br>$K_m$ = 4 [$\mu$M] | Modified from [27551076] | Cofilin activation |
| rr21 | -Lig/$\tau$ | $\tau$ = 80 [s] | This work | Ligand decay |

**Table S4. Governing equations for the species**

| Reaction | Equation | Units | Ref |
|---|---|---|---|
| rLig | rr21 | [mol/(m$^3$ s)] | This work |
| rCa_cyto | 0 | [mol/(m$^3$ s)] | This work |
| CafluxOnToPM | rr8+rr11 | [mol/(m$^2$ s)] | [17693463] |
| rReceptor | -rr1-rr2 | [mol/(m$^2$ s)] | [17693463] |
| rIP3 | -rr16 | [mol/(m$^3$ s)] | [17693463] |
| IP3fluxOnToPM | -rr14-rr15 | [mol/(m$^2$ s)] | [17693463] |
| rL_R_GD | rr3+rr4-rr5-rr6 | [mol/(m$^2$ s)] | [17693463] |
| rIP3_deg | rr16 | [mol/(m$^3$ s)] | [17693463] |
| LigfluxOnToPM | rr1+rr4 | [mol/(m$^2$ s)] | [17693463] |
| rR_GD | rr2-rr5 | [mol/(m$^2$ s)] | [17693463] |
| rGT | rr6-rr87-rr9-rr10 | [mol/(m$^2$ s)] | [17693463] |
| rGD | -rr2-rr3+rr7+rr12+rr13 | [mol/(m$^2$ s)] | [17693463] |
| rPLC | -rr8-rr9+rr13 | [mol/(m$^2$ s)] | [17693463] |
| rPLC_GT | rr9-rr11-rr13 | [mol/(m$^2$ s)] | [17693463] |
| rPIP2 | -rr14-rr15 | [mol/(m$^2$ s)] | [17693463] |
| rLR | rr1-rr3+rr6 | [mol/(m$^2$ s)] | [17693463] |
| rPLC_Ca | rr8-rr10+rr12 | [mol/(m$^2$ s)] | [17693463] |
| rPLC_Ca_GT | rr10+rr11-rr12 | [mol/(m$^2$ s)] | [17693463] |
| rL_R_GD_P | rr5 | [mol/(m$^2$ s)] | [17693463] |
| rCa_ER | 0 | | This work |
| CafluxInToER | rr17+rr18 | [mol/(m$^2$ s)] | [17483174] |
| CafluxInToEC | rr19 | [mol/(m$^2$ s)] | [17693463] |
| rCofilinAct | rr20 | [mol/(m$^3$ s)] | Modified from [24004559] |
| rCofilin | -rr20 | [mol/(m$^3$ s)] | Modified from [24004559] |

**Table S5. Geometric parameters in the spatial model**

| Cell section | Dimension | Units | Details |
|---|---|---|---|
| Neurite | Length - ~30<br>Radius - 1.25 | [$\mu$m] | |
| ER | Length - 25<br>Radius - 0.25 | [$\mu$m] | Located within the neurite |
| Nucleus | Radius - 2 | [$\mu$m] | Acts as an excluded volume |
| Soma | Radius - 5 | [$\mu$um] | |

**S3. Mechanical model**

We developed an ordinary differential equation model of actin- and tension-driven mechanics controlling neurite retraction. Signaling dynamics are coupled and feed into the mechanical model [19787029] as spatially-averaged values. We based the mechanical model framework on experimental observations and existing literature [24004559, 20159147, 16679410]. Using a framework proposed in [24004559] and experimental observations that neurite retraction as proportional to length, we developed a force balance written as:

$$\underbrace{-F_{tension}}_{\text{Force due to cortical membrane tension}} + \underbrace{F_{actin}}_{\text{Force due to actin dynamics}} = \underbrace{\frac{\eta}{k_L L}\frac{dL}{dt}}_{\text{Drag forces}} \text{ (S1),}$$

where

$$\eta = \underbrace{\mu_e}_{\text{drag coefficient for external fluid}} + \underbrace{\frac{\mu_m}{\delta}}_{\text{drag coefficient for membrane}} \text{ (S2).}$$

We arranging (Eq. S1), we find that the rate of neurite retraction could be given as

$$\frac{dL}{dt} = k_L \cdot L \cdot \frac{-F_{tension} + F_{actin}}{\eta},$$

where

$$\eta = \mu_e + \frac{\mu_m}{\delta}$$

is the drag due to both the external fluid and membrane [24004559]. $F_{tension}$ and $F_{actin}$ have dependency on signaling input. In particular, $F_{tension}$ is a function of tension, $\tau$, which itself depends on $PIP_2$. $F_{actin}$ is dependent on activated cofilin, and set amounts of total cofilin, drebrin, and actin. We can summarize the signaling inputs as $\tau = f(PIP2)$ and $F_{actin} = g(cofilinAct)$.

**S3.1 Force due to tension**

The tension force is directly proportional to the tension and is given by

$$F_{tension} = -2 \cdot \pi \cdot rTube \cdot \tau \cdot k_{Ften},$$

where $\tau$ is tension [24004559]. Tension can be thought of as a Lagrange multiplier for conserved area, and tension change has three driving components regarding area change. We construct the model so that a decrease in area in the neurite, or an increased difference in area change between the neurite and base of the neurite causes an increase in tension. The rate of area change is given by

$$\frac{dArea}{dt} = \Delta Area_{endosome} + \Delta Area_{PIP_2} \cdot f(PIP_2) + \Delta Area_{tension},$$

Where $\Delta$Area represents the rate of change in area due to each contribution. Area change is due to the internalization of activated and phosphorylated GPCRs in endosomes, the decrease in surface area due to $PIP_2$ hydrolysis that leaves the smaller DAG molecule in the membrane, and the increase in tension.

**S3.2 Area change due to endocytosis of GPCRs**

When GPCRs are both bound to ligand and phosphorylated, they are endocytosed in endosomes. This endocytosis process removes membrane and thus reduces the surface area. This rate of membrane area change is given by

$$\Delta Area_{endosome} = -\frac{d[L-R-GD-P]}{dt} \cdot 4 \cdot \pi \cdot (r_{endo})^2 \cdot \text{convertToMolecules} \cdot rTube \cdot L \cdot k_{areaEndo},$$

where $r_{endo}$ is the radius of a typical endosome.

**S3.3 Area change due to $PIP_2$ hydrolysis**

During $PIP_2$ hydrolysis, $PIP_2$ is converted into a cytosolic component, $IP_3$, and a membrane bound component, DAG. DAG is smaller than $PIP_2$, leading to a reduction in membrane surface area. This rate of membrane area change is given by

$$\Delta Area_{PIP_2} = \frac{dPIP_2}{dt}_{\text{maxInNeurite}} \cdot \delta_{AreaPIP_2} \cdot \text{convertToMolecules} \cdot rTube \cdot L \cdot k_{areaPIP_2},$$

where $\delta_{AreaPIP_2}$ is the area difference between $PIP_2$ size and DAG size. This area change is dependent on the rate of $PIP_2$ hydrolysis. $PIP_2$ dynamics have also been linked to effects on cytoskeleton, supporting the coupling of our model [12221130]. Note that in the total area change equation in S3.1, the $PIP_2$ area term is multiplied by $f(PIP_2)$ which is a sigmoidal function dependent on the $PIP_2$ density and is given by

$$fPIP2 = \frac{1}{1 + exp(-fscale(Ptrigger - PIP2_{maxinneurite}))}.$$

The sigmoidal form is based on [29473547].

**S3.4 Area change due to tension**

There is also a membrane area change due to tension. This term serves as feedback into the tension force equation because as tension changes due to area change, it can affect the rate of area change itself. This rate of membrane area change is based on [28648660] and is given by

$$\Delta Area_{tension} = 2 \cdot \pi \cdot rTube \cdot \frac{dL}{dt} \cdot \frac{\tau}{\tau_0} \cdot k_{tau}.$$

$\tau_0$ is the initial tension value.

Combining these effects, we get the follow equation for tension,

$$\frac{d\tau}{dt} = -\frac{dArea}{dt} \cdot F_{perPIP2} \cdot \frac{1}{A0} \cdot k_{tension}.$$

**S3.5 Actin force**

The actin force is influenced by various actin-related proteins. We note that actin dynamics are highly complicated quite extensively studied [24209839, 10.1523/JNEUROSCI.11-07-01918.1991, 10.1016/0896-6273(88)90124-9, 20159147]. We also highlight that we experimentally observe unusual actin dynamics during this neurite retraction. Therefore, we present a phenomenological model for actin dynamics based on a stress-strain framework. We construct that model such that cofilin reduces the actin pushing force, and drebrin and actin slow the reduction in actin pushing force. Eventually the decrease in pushing force becomes a negative force that can be attributed to retrograde flow [10.1016/S0955-0674(97)80152-4, 24198333]. Therefore, actin force is given by

$$\frac{dF_{actin}}{dt} = F_{perActin} \cdot \frac{dL}{dt} \cdot \frac{k_{cofilin} + k_{actin} + k_{drebrin} + k_{const}}{L_0},$$

where the k contributions are from cofilin [10.1002/(SICI)1097-0169(1998)39:2<172::AID-CM8>3.0.CO;2-8], actin [10.1523/JNEUROSCI.11-07-01918.1991], and drebrin [21175132,23696644]. The k equations are given by

$$k_{cofilin} = gCofilinAct \cdot cofilinIC/(1[uM]),$$

$$k_{actin} = -0.5actinIC/10, \text{ and}$$

$$k_{drebrin} = -0.5drebrinIC/4.$$

The cofilin contribution is again dependent on a sigmoidal function [29473547] that is dependent on activated cofilin (cofilinAct), and is given by

$$gCofilinAct = \frac{1}{1 + exp(-gscale(CofTrigger - (CTotal - cofilinAct_{avgincyto})))}.$$

The various parameters in the mechanical model are given in Table S6.

**Table S6. Parameters in the mechanical model**

| Reaction | Equation | Units | Ref | Description |
|---|---|---|---|---|
| $k_L$ | 3.2 x $10^6$ | [1/$m^2$] | This work | scaling |
| $\mu_e$ | 8.9 x$10^{-4}$ | [Pa s] | [24004559] | Friction fluid |
| $\mu_m$ | $10^{-9}$ | [N s/m] | [24004559] | Membrane friction |
| $\delta$ | 5 | [nm] | [24004559] | Bilayer thickness |
| rHead | 5 | [$\mu$m] | Geometric constraint | Soma radius |
| rTube | 1.25 | [$\mu$m] | Geometric constraint | Neurite radius |
| $L_0$ | 30 | [$\mu$m] | Geometric constraint | Neurite length |
| $k_{Ften}$ | 0.5x$10^{-4}$ | unitless | This work | scaling |
| $\delta_{AreaPIP_2}$ | 30x$10^{-20}$ | [$m^2$] | Inspired by [10.1146/annurev. biophys.31.08290 1.134259] | Change in area from PIP2 to DAG |
| convertToMolecules | 6.022 x $10^{23}$ | [1/mole] | Avogadro's number | |
| $k_{areaPIP2}$ | 1.75x$10^2$ | unitless | This work | scaling |
| $(r_{endo})^2$ | 2.5x$10^{-15}$ | [$m^2$] | [31017840] | Radius squared of an endosome |
| $k_{areaEndo}$ | 1.2 | unitless | This work | scaling |
| A0 | 2.2444x$10^{-10}$ | [$m^2$] | Geometric constraint | Area of neurite membrane |
| $k_{tension}$ | 75 | unitless | This work | scaling |
| $F_{perPIP2}$ | 10 | [pN/nm] | This work | Force per PIP2 |
| Fscale | 4x$10^9$ | [$m^2$/mol] | This work | scaling |
| Ptrigger | 1.5x$10^{-9}$ | [mol/$m^2$] | This work | PIP2 trigger |
| $F_{peractin}$ | 40 | [pN] | This work | Force per actin |
| gscale | 9 | [1/$\mu$M] | This work | scaling |

| | | | | |
|---|---|---|---|---|
| CofTrigger | 1.3 | [$\mu$M] | This work | CofilinAct trigger |
| CTotal | 2 | [$\mu$M] | This work | Total cofilin |
| $k_{tau}$ | 0.1 | unitless | This work | Scaling |
| $\tau_0$ | 254.6 | [pN/nm] | This work – determined; inspired by [20159147] | Initial tension |
| $k_{const}$ | 2.375 | Unitless | This work - determined | |
| $Factin_0$ | 100 | [pN] | This work – inspired by [20159147] | Initial $F_{actin}$ |

**S3.6 Overexpression trials**

To capture protein overexpression experiments for actin, cofilin, and drebrin, the concentration of overexpressed protein was input into the corresponding protein's k equation. For example, actin is believed to be overexpressed from about 10 $\mu$M to 30 $\mu$M, so actinIC goes from 10 to 30 in actin's k equation. In this way, we capture the effects of protein overexpression on actin force dynamics. See Table S7 for the estimated protein concentrations during each overexpression trial.

Table S7. Actin-related protein concentrations

| Protein | Original concentration [$\mu$M] | Overexpressed concentration [$\mu$M] | Reference |
|---|---|---|---|
| Actin | 10 | 30 | [27551076] and this work |
| Cofilin | 2 | 12 | [27551076] and experimental observation from this work |
| Drebrin | 1 | 4 | [10.1046/j.1471-4159.1996.66030980.x] |

**S3.7 Hyperosmotic stress model**

To elicit the role of tension in neurite retraction dynamics, hyperosmotic stress experiments were conducted. During these trials, neurite retraction was observed in the absence of ligand stimulation, confirming the role of tension in retraction. To model hyperosmotic conditions, we assumed that that hyperosmotic environment will trigger volume reduction and an initial increase in membrane tension. Therefore, we increase our initial tension value in the absence of ligand input.

**S4 Western Blots to assess protein levels used in modeling**

**Actin**

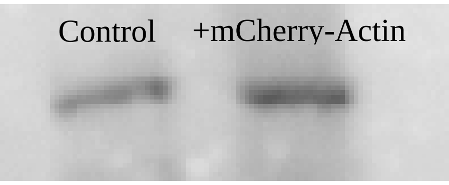


**Cofilin**

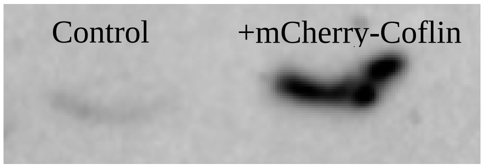


**V1-2 (attached) Videos showing neurite retraction with carbachol stimulation seen for PC12 cells labeled with Calcium Green (left) and DIC (right). The corresponding screen shots are shown in Fig 1 and below. Experimental details can be found in the text.**

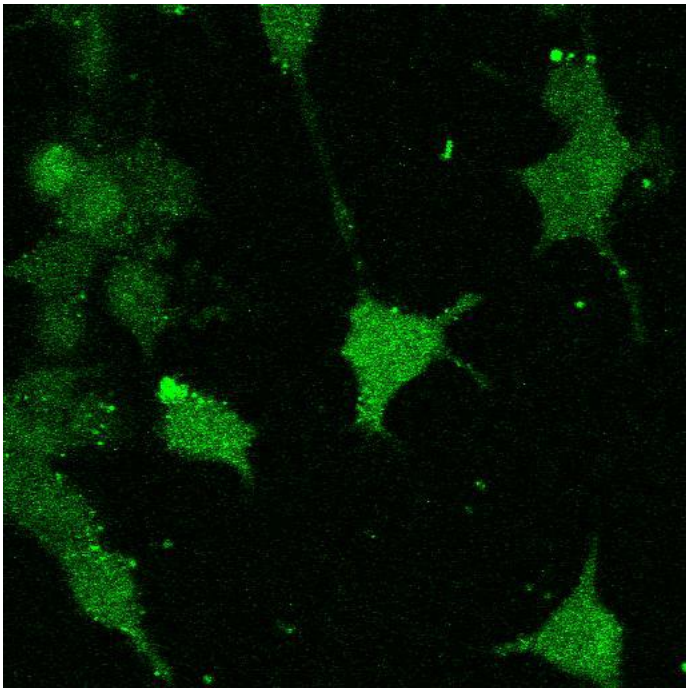

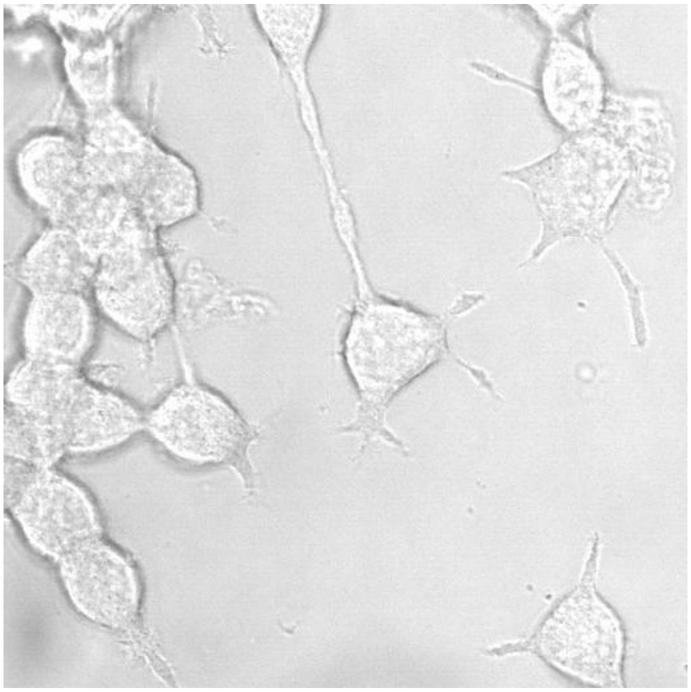

**V3 (attached) Neurite retraction with carbachol stimulation as monitored by eCFP-B2R where the associated screen shots are shown below and can be found in Fig2 B-C. Experimental details can be found in the text.**

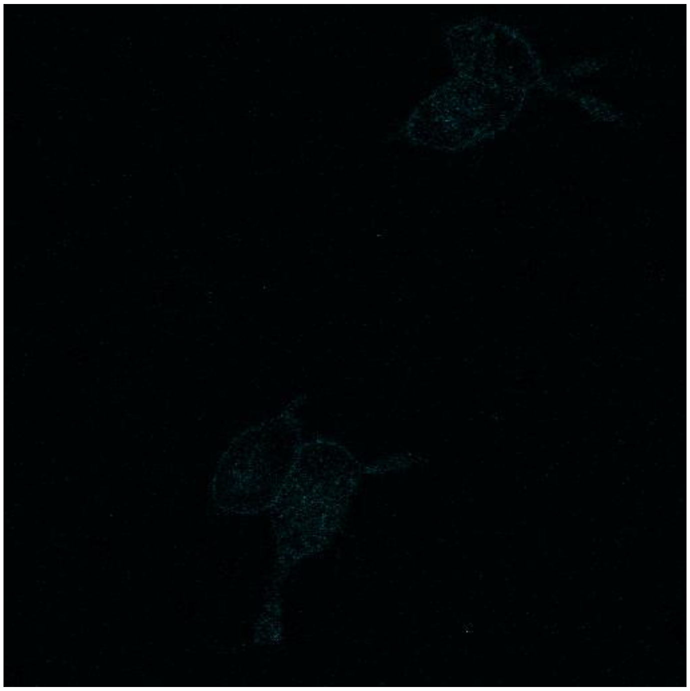

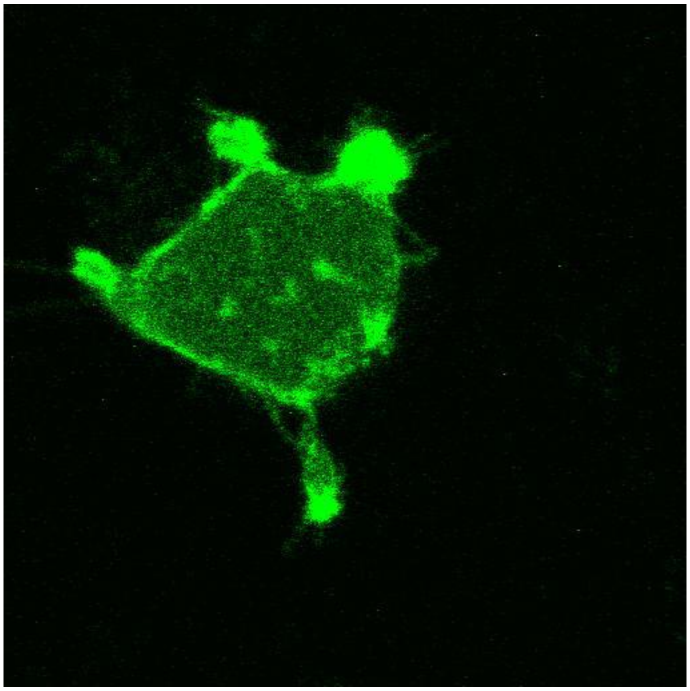

**V4 (attached) The behavior of PI(4,5)$P_2$ with carbachol-induced neurite retraction as followed by PH-PLCδ1 where the screen shots are shown below and in Fig2 F-G. Experimental details can be found in the text.**

**V5 (attached) The behavior of mCherry-actin with carbachol-induced neurite retraction where the screen shots are shown below and in Fig. 4 D, G, J. Experimental details can be found in the text.**

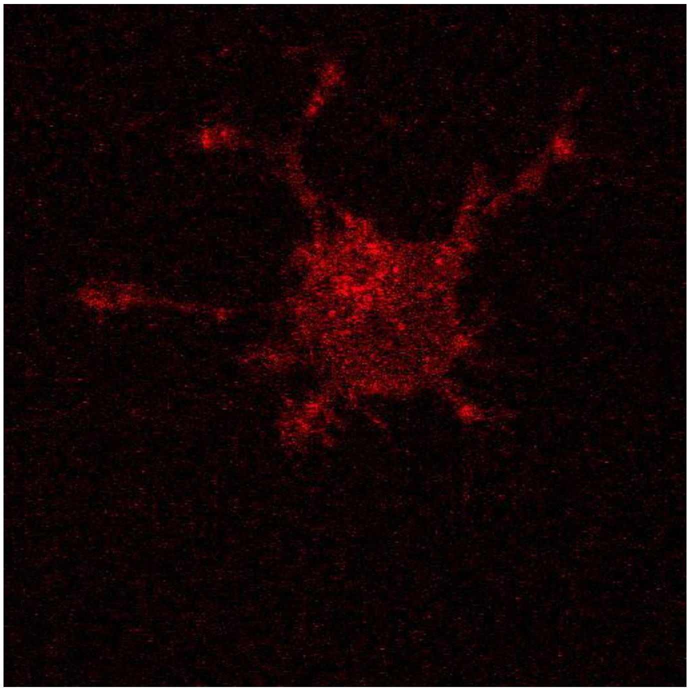

**V6 (attached) The behavior of mCherry-cofilin with carbachol-induced neurite where the screen shots are shown below and in Fig4 E, H, K. Experimental details can be found in the text.**

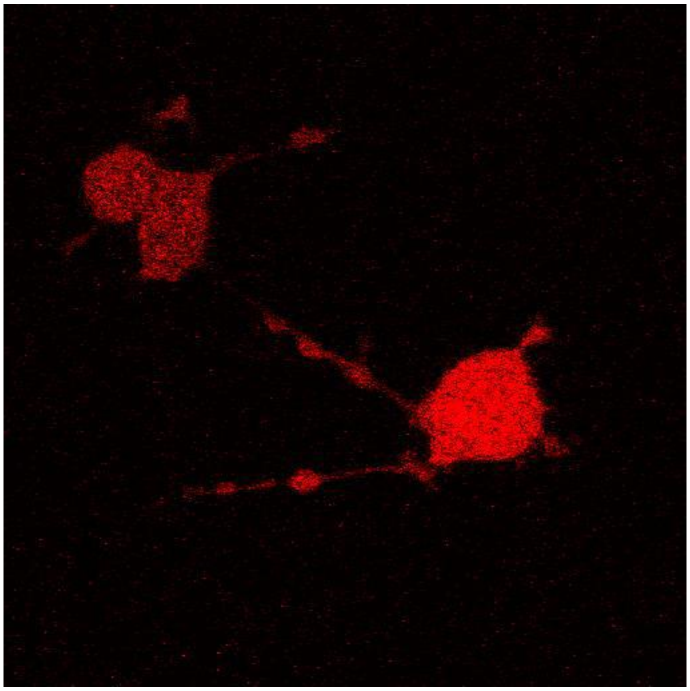
